# Altered DNA repair pathway engagement by engineered CRISPR-Cas9 nucleases

**DOI:** 10.1101/2022.03.10.483793

**Authors:** Vikash P. Chauhan, Phillip A. Sharp, Robert Langer

## Abstract

CRISPR-Cas9 introduces targeted DNA breaks that engage competing DNA repair pathways, producing a spectrum of imprecise insertion/deletion mutations (indels) and precise templated mutations (precise edits). The relative frequencies of these pathways are thought to primarily depend on genomic sequence and cell state contexts, limiting control over mutational outcomes. Here we report that engineered Cas9 nucleases that create different DNA break structures engage competing repair pathways at dramatically altered frequencies. We accordingly designed a Cas9 variant (vCas9) that produces breaks which suppress otherwise dominant nonhomologous end-joining (NHEJ) repair. Instead, breaks created by vCas9 are predominantly repaired by pathways utilizing homologous sequences, specifically microhomology-mediated end-joining (MMEJ) and homology-directed repair (HDR). Consequently, vCas9 enables efficient precise editing through HDR or MMEJ while suppressing indels caused by NHEJ in dividing and non-dividing cells. These findings establish a new paradigm of targeted nucleases custom-designed for specific mutational applications.

**Teaser:** CRISPR-Cas9 can be designed to make otherwise infrequent precise editing pathways dominant in dividing and non-dividing cells

## Introduction

Genome editing with CRISPR-Cas9 harnesses cellular double-strand break (DSB) repair pathways, such as non-homologous end-joining (NHEJ), microhomology-mediated end-joining (MMEJ), and homology-directed repair (HDR), to mutate targeted loci (*1, 2*). NHEJ produces semi-random insertion/deletion mutations (indels) that are generally small. In contrast, MMEJ makes sequence-specific indels that can be small or large by using small homologous sequences flanking the DSB. Meanwhile, HDR utilizes a homologous DNA repair template that recombines with the DSB site to create specific mutations called precise edits. These repair pathways compete to repair each DSB through multi-faceted regulation (*3–7*), though NHEJ is the dominant mechanism (*8, 9*). The relative frequencies of these repair pathways and the mutations they produce follow a predictable distribution that is thought to be defined by the sequence of the targeted locus and the cell state (*10, 11*). A central goal of the field is to develop means for controlling the relative frequencies of particular repair pathways, primarily with the aim of making precise editing by HDR or specific indels by MMEJ more frequent. Several approaches have sought to increase the efficiency of precise editing, including varied template design (*12–14*), template stabilization (*15*), template localization (*16*), small molecule NHEJ inhibition (*17, 18*), cell synchronization (*12*), and modulation of regulatory factors that suppress precise HDR and MMEJ (*19–22*). While useful, these strategies involve manipulation of cell state or the addition of exogenous factors, which may not be feasible in many genome editing applications.

Cas9 nucleases that are intrinsically biased toward DSB repair by one pathway or another would be highly desirable, yet a design paradigm for altering repair pathway engagement has not been established. Thus far Cas9 has been extensively engineered for enhanced on-target versus off-target DNA cutting specificity (*23, 24*) or altered protospacer-adjacent motif (PAM) recognition (*25, 26*). These two parameter spaces for Cas9 nuclease design have been well explored, and intriguingly some of these Cas9 variants display modestly different frequencies of DSB repair pathways through unknown mechanisms (*27*). Without a clear mechanism driving these repair biases, it remains unexplored whether Cas9 nucleases can be engineered to specifically alter repair outcomes. Separately, it has been established that DNA break structure influences repair mechanisms (*28, 29*). Utilizing Cas9 nucleases to make blunt cut DSBs preferentially leads to indels through NHEJ, whereas producing large 5’ overhangs by generating offset nicks with Cas9 nickases instead promotes DNA resection, MMEJ, and HDR in different contexts (*29, 30*). Intriguingly, natural and engineered Cas9 variants produce slightly varied DSB structures following cutting, ranging from blunt cutting to occasionally producing staggered cuts with 1bp overhangs (*31, 32*). We hypothesized that Cas9 could be engineered to produce larger staggered DSBs that would correspondingly bias repair pathway engagement. Here we explore this potential design paradigm for altering repair outcomes for Cas9 nucleases.

## Results

We first investigated whether Cas9 could be engineered to produce different DNA break structures that bias repair outcomes. We reasoned that mutations in residues located at the DNA substrate interface of Cas9 could modulate positioning of the DNA cuts (Fig. 1A). To test this, we selected residues in *S. pyogenes* Cas9 with known or proposed interactions with the substrate strands (*23, 24, 33*). These fourteen residues were either basic or polar and we mutated each to a hydrophobic Alanine to reduce putative DNA interactions (Fig. 1B and fig. S1A-E). To assess break structures, we applied a well-characterized approach in which two concurrent DSBs are created and the resulting DNA end junctions are analyzed for sequences resulting from blunt versus staggered cutting (fig. S2A) (*31*). We thereby determined break structures for these engineered variants by making two nearby DSBs at the *EMX1* locus in HEK293T cells and analyzing junction sequences following deep sequencing of the amplified locus (fig. S2B). We concomitantly measured precise editing and indel frequency for these variants by combining them with HDR templates that produce small sequence replacements at the *EMX1* and *CXCR4* loci in HEK293T cells and analyzing alterations using deep sequencing of the amplified locus (Fig. 1C). We found that wild-type Cas9 almost uniformly produced blunt cuts, whereas several Cas9 single-mutant variants induced staggered cuts from 2-6bp (Fig. 1D). We then examined the relative frequencies of different repair pathways engaged by these engineered Cas9 variants. Several of the Cas9 single-mutant variants reduced the frequency of indels while increasing precise editing frequency (Fig. 1E and fig. S3A). There was a strong correlation between precise editing frequency and staggered cutting for these Cas9 variants, indicating that DNA break structure regulated repair pathway outcomes (Fig. 1F).

**Figure 1.**
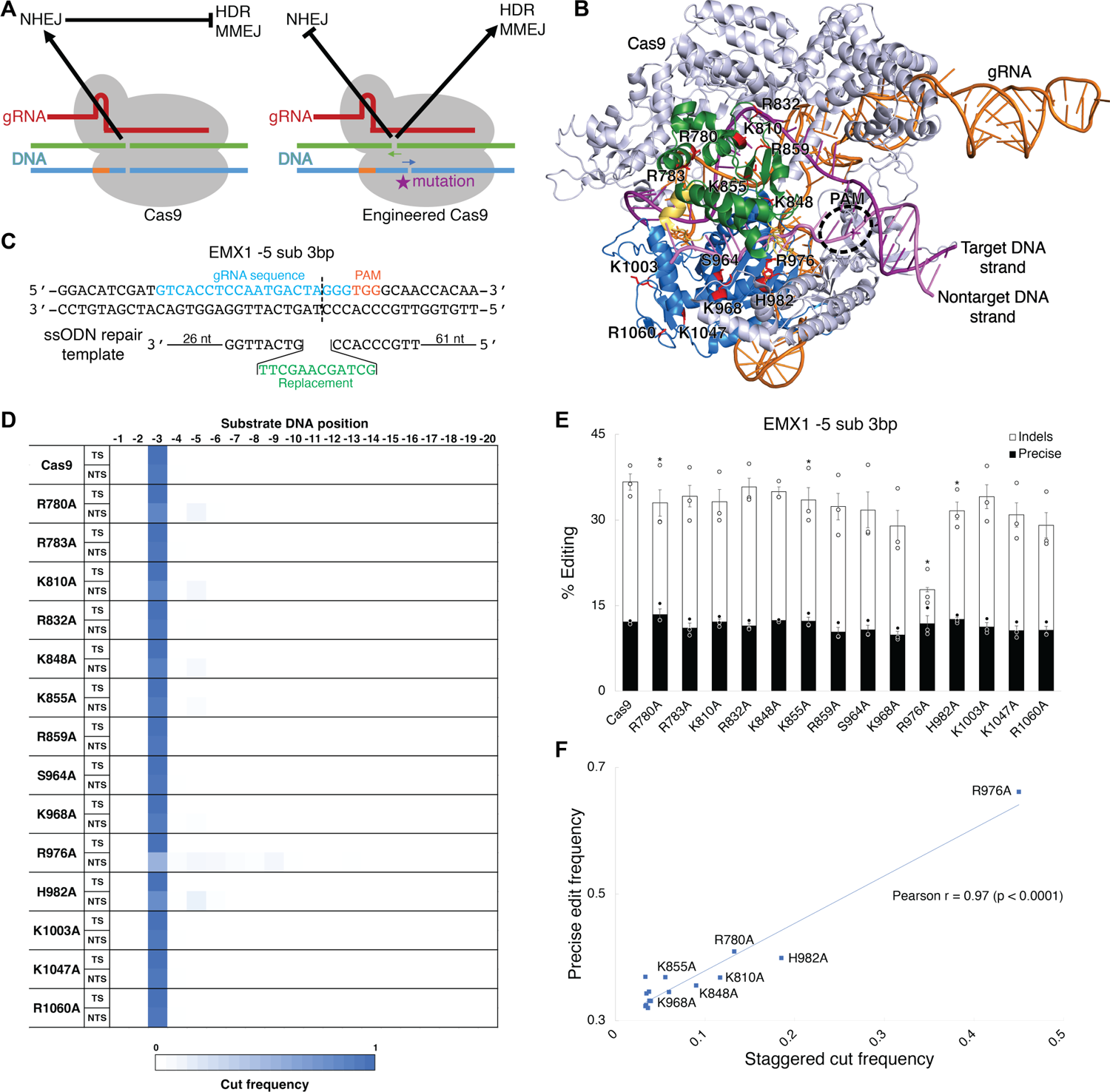
Engineered Cas9 variants that make staggered cuts have altered repair outcomes. (**A**) Model of the balance between NHEJ and HDR or MMEJ repair pathways for Cas9 variants. (**B**) Cas9 residues at the interface with the substrate DNA strands that were selected for Alanine substitution and altered repair pathway screening. The mutated residues (red) are located in either the mobile HNH domain (green) or the immobile RuvC domain (blue), which are connected by linker segments (yellow). (**C**) Design of a gRNA and HDR template to introduce precise edits. (**D**) DNA break positions in the target strand (TS) and non-target strand (NTS) for engineered Cas9 single-mutant variants. (**E**) Screen of precise editing and indel frequencies for engineered Cas9 single-mutant variants using an HDR template. (**F**) Correlation between staggered cut frequency and precise editing frequency across Cas9 variants from the Alanine-substitution screen. * indicates p < 0.05 for precise editing frequency compared to wild-type Cas9. Data were analyzed by deep sequencing and represent means of *n* = 3 independent replicates with standard errors.

We subsequently explored whether Cas9 mutations that modulate DNA break structure could be utilized to engineer Cas9 for enhanced precision. We combined mutant residues that changed repair outcomes for the single-mutant variants and identified several mutant combinations that significantly increased precise editing frequency and reduced indel frequency relative to wild-type Cas9 (Fig. 2A,B and fig. S3B). We tested these engineered Cas9 variants by applying HDR to convert a green fluorescent protein (GFP) transgene inserted in the genome of HEK293T cells to blue fluorescent protein (BFP) (*17*). One variant, R976A-K1003A, most effectively increased precise editing frequency while decreasing indel frequency (fig. S3C-E). Yet while this variant greatly altered the frequency of these competing repair pathways, its total editing activity was lower than that of wild-type Cas9 at all loci tested. To address the challenge of lower total editing activity, we investigated whether the lost total editing activity could be rescued by additional mutations. The R976A mutation produced the greatest change in repair pathway frequency along with the largest decrease in total editing activity and sits at one end of the nontarget cleft near where the substrate DNA strands separate (Fig. 1E). We thought that R976 might regulate repair pathway frequency by affecting how the nontarget strand sits in its cleft and where it is cut, while also controlling total activity by aiding in substrate strand separation prior to cutting. To test whether mutating residues nearby A976 to Arginine could rescue activity without reverting changes to repair pathway frequency, we mutated six residues in the R976A-K1003A variant to produce a set of triple- and quadruple-mutants (Fig. 2C). We assessed these new variants using a similar HDR assay at the *VEGFA* and *EMX1* loci in HEK293T cells and analyzed using deep sequencing (Fig. 2A). This led to one variant, S55R-R976A-K1003A-T1314R, that maximally rescued activity (Fig. 2D,E and fig. S3F,G). Importantly, this variant did not increase non-specific genome activity but surprisingly improved on-target versus off-target editing specificity compared to wild-type Cas9 (fig. S4). Since Cas9^S55R-R976A-K1003A-T1314R^ was the most intriguing and active variant of Cas9, we named it vCas9.

**Figure 2.**
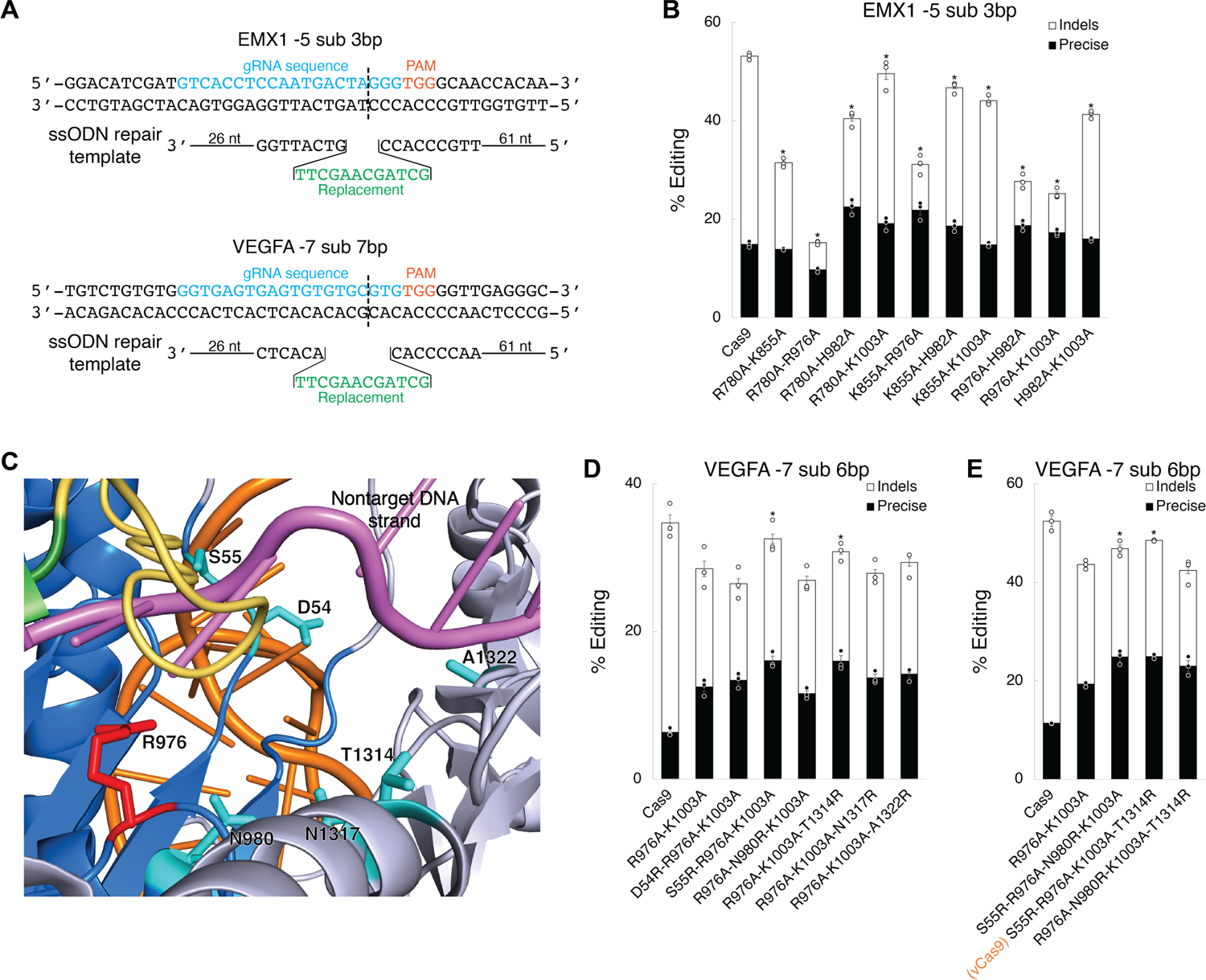
Design of Cas9 variants with altered repair pathway frequencies. (**A**) Design of gRNAs and HDR templates to introduce precise edits. (**B**) Screen of precise editing and indel frequencies for engineered Cas9 double-mutant variants using an HDR template. (**C**) Cas9 residues proximal to R976 that were selected for Arginine substitution and activity rescue screening. The mutated residues (teal) are located near the nontarget substrate DNA strand (pink). (**D**, **E**) Screen of precise editing and indel frequencies for engineered Cas9 (**D**) triple-mutant and (**E**) quadruple-mutant variants using an HDR template. * indicates p < 0.05 for precise editing frequency compared to wild-type Cas9 in **B** or for total editing frequency compared to R976A-K1003A in **D**, **E**. Data were analyzed by deep sequencing and represent means of *n* = 3 independent replicates with standard errors.

To clarify the mechanisms by which the above variants altered DNA repair pathway frequency, we further studied DNA break structure and editing outcomes. We found that vCas9 showed a large shift in break structure by predominantly making staggered cuts of 6bp or larger (Fig. 3A). To determine whether this break structure difference affected DNA resection and repair pathway frequency, we then analyzed indel distributions and mechanisms for wild-type Cas9 and vCas9 at several loci in HEK293T cells using deep sequencing (Fig. 3B and fig. S5A-D). We found that vCas9 induced indels of larger size than those by wild-type Cas9 (Fig. 3C) and greatly suppressed NHEJ deletions and insertions while promoting MMEJ deletions (Fig. 3D and fig. S6A,B). Specifically, vCas9 shifted indels from NHEJ repair to MMEJ repair utilizing larger microhomologies, indicating enhanced DNA end resection or strand separation following the staggered DNA cuts (fig. S5E). In the presence of an HDR template, vCas9 consistently made HDR dominant, with the remaining minor repair outcomes comprised mostly of NHEJ insertions and MMEJ deletions (Fig. 3E and fig. S5F). We also assessed wild-type Cas9 and vCas9 repair frequencies following treatment of HEK293T cells with inhibitors of MMEJ (Rucaparib) or NHEJ (NU7026). Inhibiting NHEJ with wild-type Cas9 led to editing patterns that resembled vCas9, while inhibiting MMEJ largely ablated editing by vCas9 but not wild-type Cas9 (fig. S5G,H). We further contrasted indel distributions without or with an HDR template at several loci in HEK293T cells to understand how break structure impacts competition between repair pathways. Remarkably, the degree to which precise editing outcompetes an indel was strongly correlated with indel size, and vCas9 increased the degree to which repair using homologous sequences outcompeted any indel (Fig. 3F). Together, these data demonstrate that vCas9 suppressed NHEJ in favor of repair pathways that use homologous sequences, hence promoting HDR or MMEJ.

**Figure 3.**
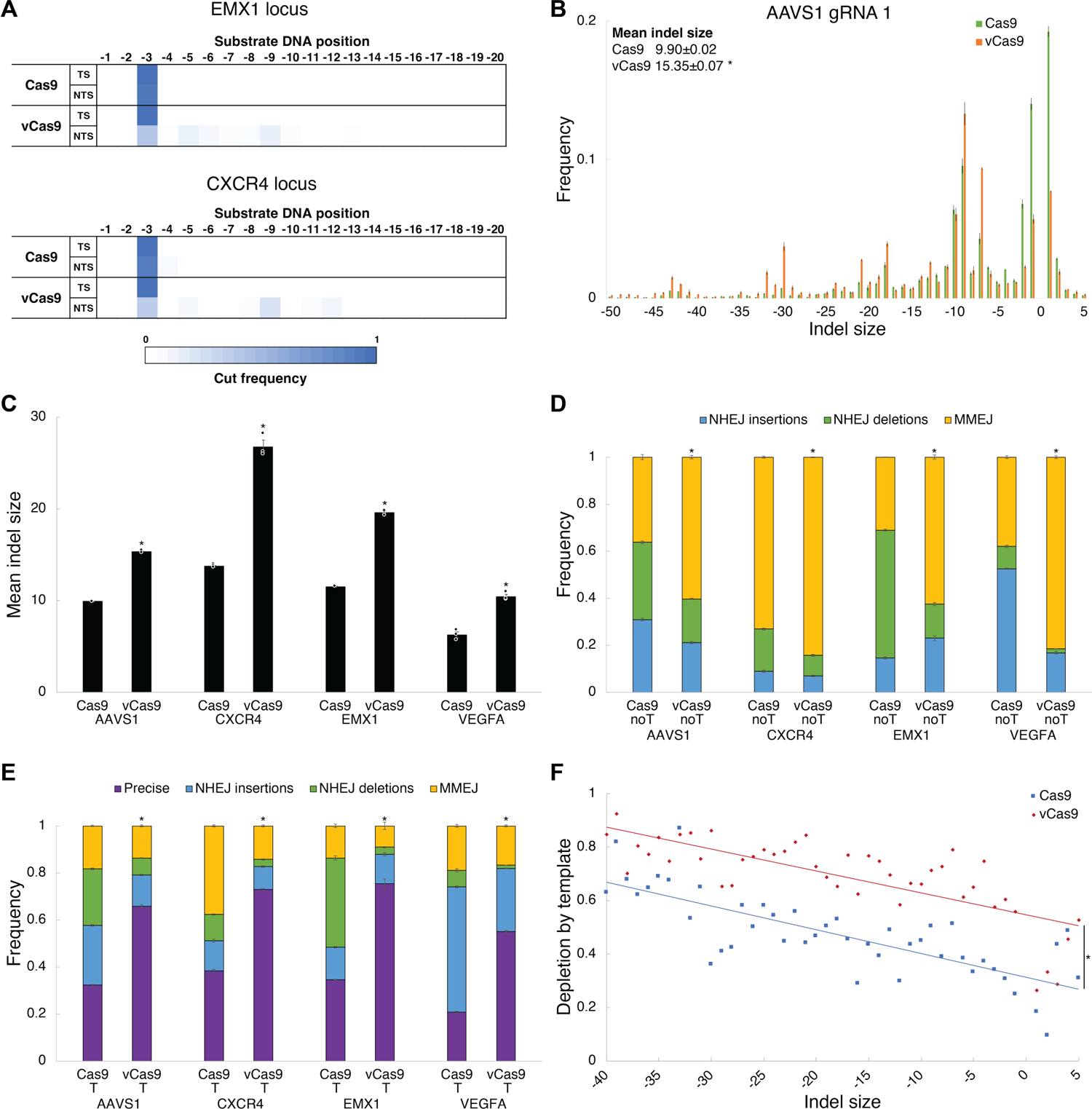
Staggered cutting by vCas9 suppresses NHEJ and makes MMEJ or HDR dominant. (**A**) DNA break positions in the target strand (TS) and non-target strand (NTS) for engineered Cas9 variants. (**B**) Distributions of indel sizes induced. (**C**) Mean indel size induced at several loci. (**D**, **E**) Frequencies of repair pathways engaged (**D**) without (noT) and (**E**) with (T) a repair template at several loci. (**F**) Degree of depletion by a repair template for indels of different sizes averaged over several loci. The fraction depleted is increased by vCas9 at all indel sizes. * indicates p < 0.05 for indel size compared to wild-type Cas9 in **B**, **C**, for NHEJ frequency compared to wild-type Cas9 in **D**, **E**, or for indel depletion compared to wild-type Cas9 in **F**. Data were analyzed by deep sequencing and represent means of *n* = 3 independent replicates with standard errors.

We next explored the robustness of these effects on DNA repair pathway frequency with respect to diverse editing contexts. To clarify sequence dependence, we assessed editing at several loci in HEK293T cells using HDR templates producing small edits with analysis by deep sequencing. Reassuringly, vCas9 significantly and consistently altered DNA repair pathway frequency at all loci (Fig. 4A). Whereas Cas9 demonstrated precise editing frequencies of 9.9-37.5% (mean 24.1%), vCas9 increased these to 43.3-73.7% (mean 58.3%), corresponding to a 1.4- to 2.8-fold (mean 1.9-fold) suppression of indel frequency. We also assessed whether vCas9-produced edits are precise and functional by again applying HDR to convert a GFP transgene to BFP. Indeed, vCas9 greatly increased precise gene conversion while suppressing indel frequency (Fig. 4B,C). To examine cell-type dependence, we applied similar HDR templates to produce small edits at several loci in HeLa, A549, and Panc1 cells. In each cell model and locus, vCas9 consistently altered precise editing and indel frequencies (fig. S7A-C). We further tested two other precise edit types of interest, large insertions from double-stranded templates and untemplated precise collapse of duplications (Fig. 4D,E) (*11*). For both edit types, vCas9 similarly favored precise editing and suppressed indels (Fig. 4F,G). We also compared to other engineered Cas9 variants and fusions and found that repair outcomes for vCas9 are more biased toward precise editing (fig. S8A,B) (*20, 21, 27*). These data indicate that vCas9 robustly suppressed indels and promoted precise editing across varied loci, cells, and edit types.

**Figure 4.**
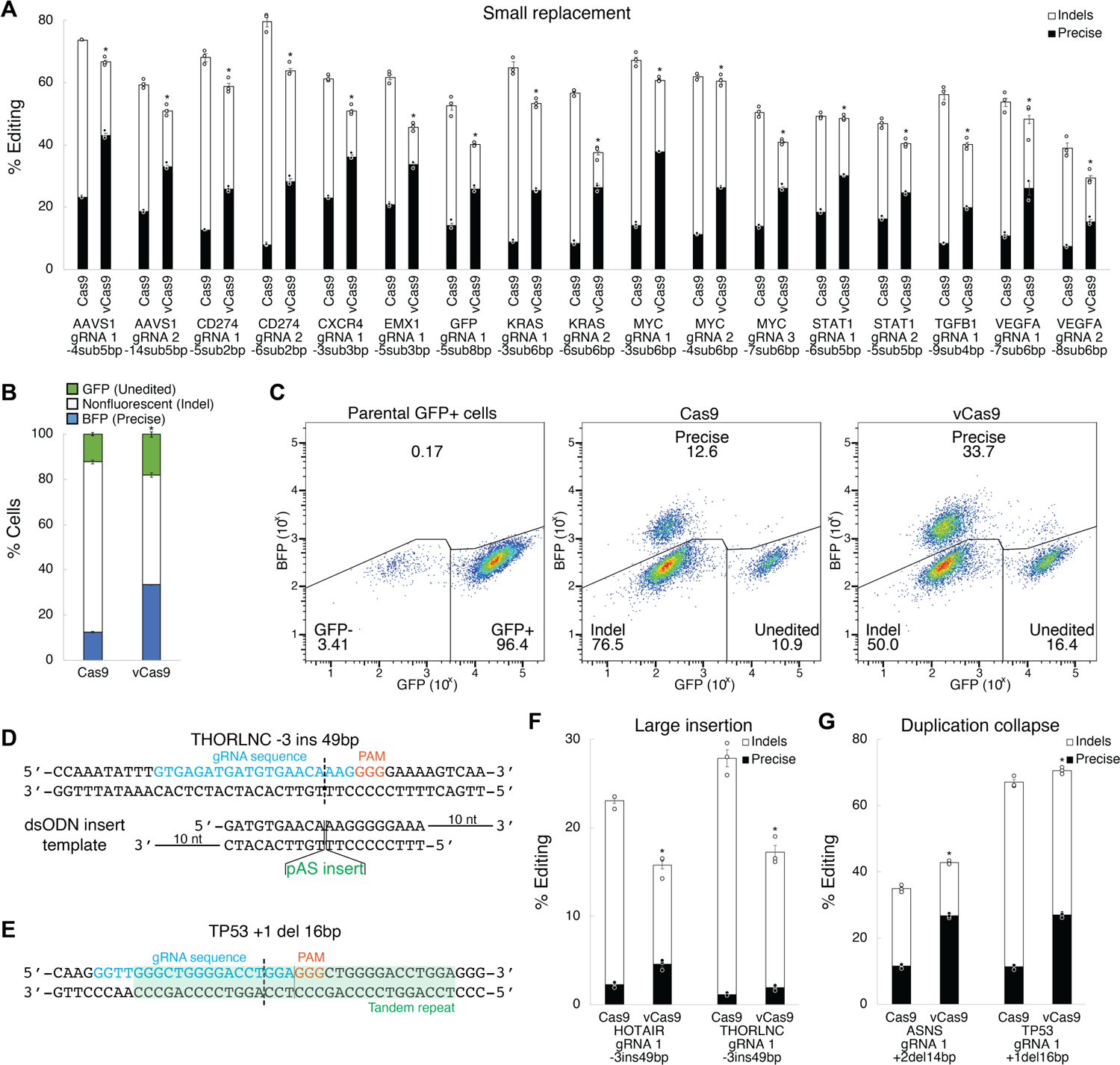
vCas9 enhances precise editing and suppresses indels across diverse editing contexts. **(A)** Precise editing and indel frequencies using small replacement (<10 bp) HDR templates at several loci. (B) Frequency of precise gene conversion from GFP to BFP. (**C**) Flow cytometry plots showing BFP-positive (indicating precise edits), nonfluorescent (indicating indels), and GFP-positive (indicating unedited cells) cell population fractions. (**D**) Design of a gRNA and HDR template to introduce a large insertion. (**E**) Design of a gRNA to induce untemplated collapse of duplications. (**F**) Precise editing and indel frequencies using large insertion (∼50 bp) templates at several loci. (**G**) Precise editing and indel frequencies for untemplated collapse of duplications (10-20 bp) at several loci. * indicates p < 0.05 for precise editing frequency compared to wild-type Cas9. Data were analyzed by deep sequencing in **A**, **F**, **G** or flow cytometry in **B**, **C**

A major limitation of CRISPR technology for precise editing to generate genetic models or treat certain diseases is the lack of HDR in non-dividing cells (*9, 19*). In contrast, MMEJ and NHEJ are active in non-dividing cells (*9, 34*). To address this, we next investigated whether vCas9 might produce precise editing in non-dividing cells by developing an MMEJ-driven templated strategy. This method, termed microhomology-directed recombination (MDR), utilizes partially double-stranded DNA templates with single-stranded microhomology arms complementary to sequences distal to the DSB ends (Fig. 5A). These MDR templates putatively enable replacements of small segments of genomic DNA situated between the two microhomology arms. We tested MDR in HEK293T cells using the GFP to BFP gene conversion assay with varied MDR template designs (fig. S9A). We found that the MDR templates enabled moderately efficient precise editing, with large increases in precise editing frequency when using vCas9 as compared to wild-type Cas9 (fig. S9B,C). Under the same conditions, MDR generated a variety of precise edit types including substitutions, insertions, and deletions, as analyzed by deep next-generation sequencing (Fig. 5A and fig. S10A-E). For each type of precise edit, vCas9 greatly increased precise editing frequency (Fig. 5B). Considering these findings, we tested MDR in quiescent primary human dermal fibroblasts, an established model of G0 non-dividing cells (*35*). Following induction of quiescence in these fibroblasts over seven days (fig. S11A,B), we assessed MDR for making small precise edits (Fig. 5A and fig S10A-E). We found that these precise edits were more frequent than all other types of edits in these cells (Fig. 5C). This was probably because vCas9-generated DSBs near-completely precluded other pathways leading to domination by precise editing.

**Figure 5.**
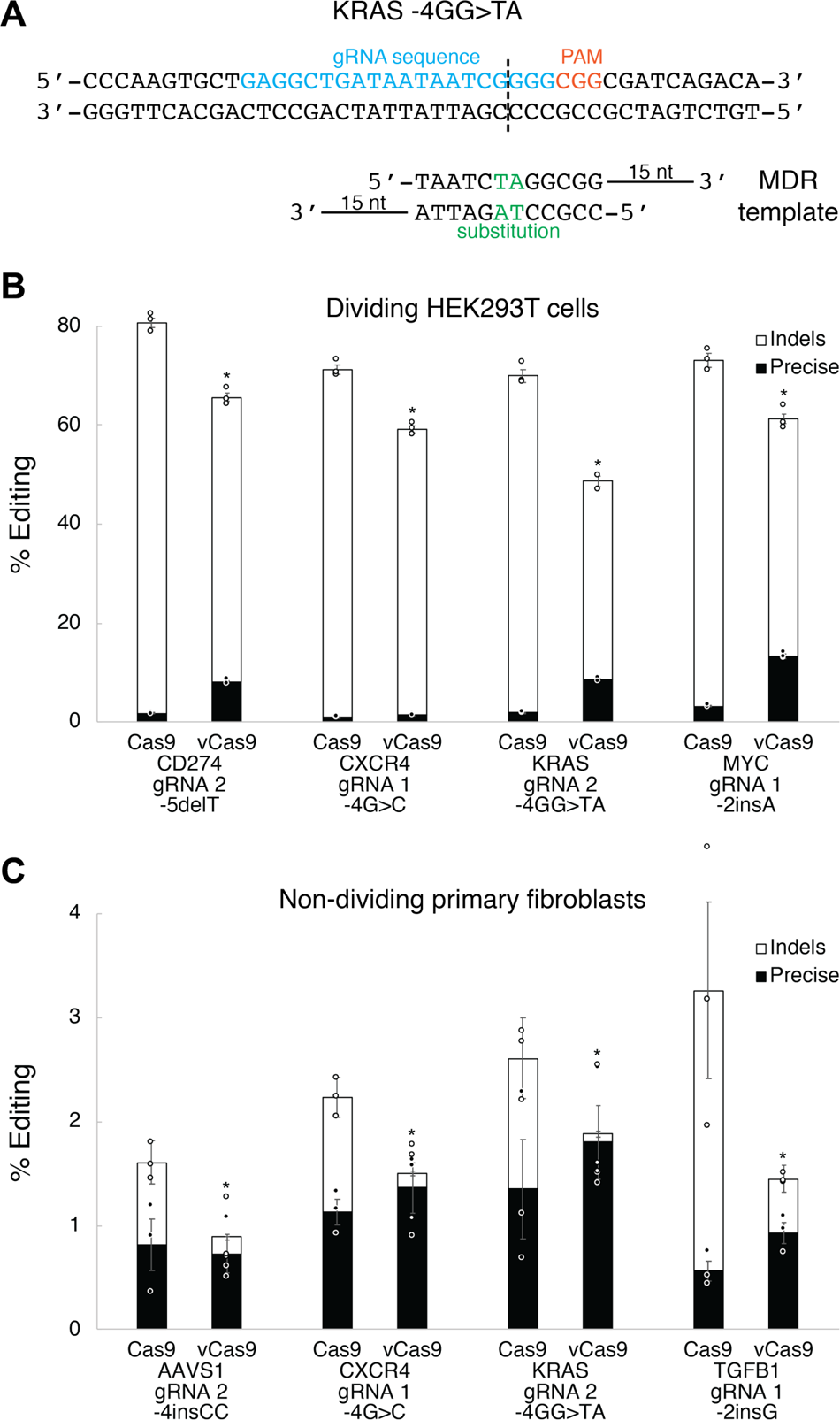
vCas9 enables efficient precise editing through MMEJ in dividing and non-dividing cells. **(A)** Design of a gRNA and MDR template to introduce a precise edit. (**B**) Precise editing and indel frequencies using MDR templates in dividing HEK293T cells. (**C**) Precise editing and indel frequencies using MDR templates in non-dividing (quiescent) primary human dermal fibroblasts. * indicates p < 0.05 for precise editing frequency compared to wild-type Cas9. Data were analyzed by deep sequencing and represent means of *n* = 3 independent replicates with standard errors.

## Discussion

Our findings support a model for how mutations in Cas9 residues at the DNA substrate interface can modulate cut positioning and downstream repair pathway outcomes. In this model, these mutations reduce protein-DNA interactions promoting movement of the DNA strands within the binding cleft (Fig. 6A,B). Notably, the mutant residues that most affected DNA break structure, R976A and H982A, are located at the 3’ end of the nontarget strand cleft near the nuclease site (fig. S1C). These mutations likely allow shifting of the nontarget DNA strand within the cleft, positioning the cut two or more nucleotides in the 5’ direction. This would produce staggered cuts that feature 5’ overhang DSBs. Processing of such 5’ overhangs would favor DNA resection and thus MMEJ or HDR repair versus NHEJ. Consistent with this model, DSBs with large 5’ overhangs produced by offset nicks have been found to promote DNA resection, MMEJ, and HDR in different contexts (*29, 30*). Our data and model broadly indicate that repair by many types of genome editors could be influenced by mutations that affect DNA break structure.

**Figure 6.**
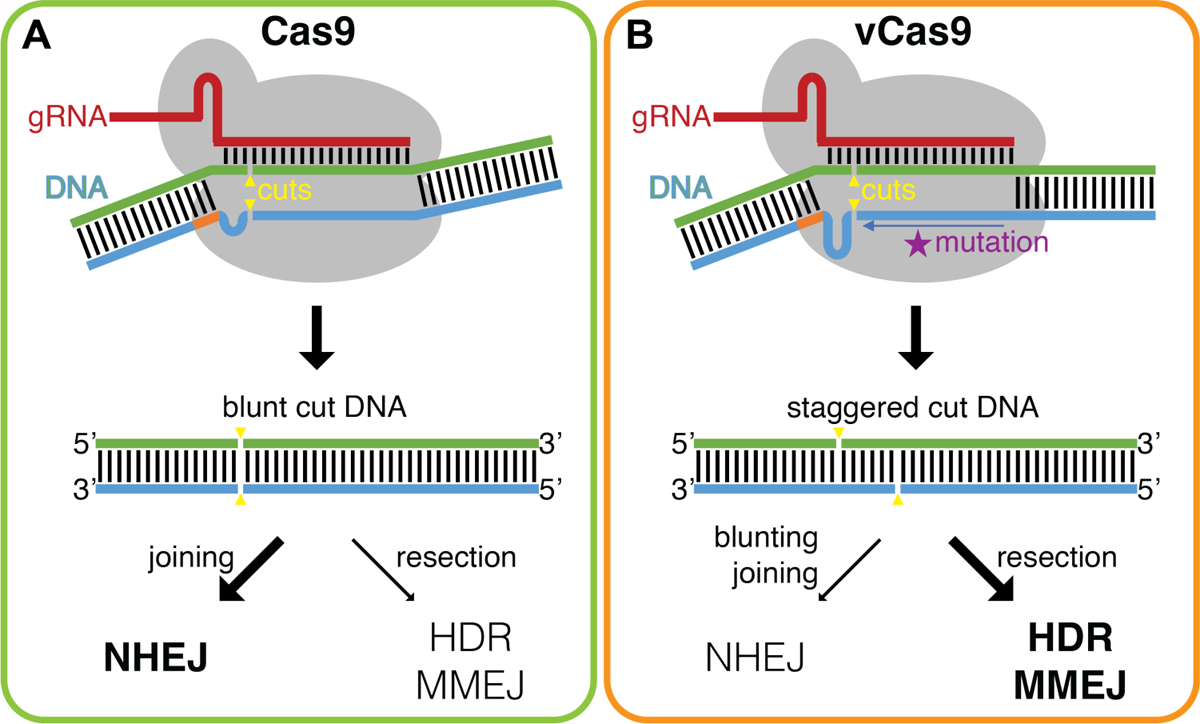
Model for how engineered Cas9 variants alter repair outcomes. (**A**) Cas9 primarily produces blunt cuts, which promote DNA end joining while limiting resection. This leads to repair dominated by NHEJ over MMEJ and HDR. (**B**) Cas9 variants like vCas9 create staggered cuts, which promote DNA resection and inhibit end joining. This enhances repair by MMEJ and HDR while suppressing NHEJ.

To our knowledge, this is the first demonstration that CRISPR-Cas9 genome editors can be engineered to intentionally alter their propensity for different pathways of DNA repair. We have shown that vCas9 suppresses NHEJ and promotes pathways that enable precise editing, such as HDR and MMEJ. These changes are not subtle, with a pronounced shift in the dominant repair pathway from NHEJ to HDR or MMEJ. Furthermore, we have shown that vCas9 makes efficient precise edits in non-dividing cells by MMEJ. These properties suggest that variants like vCas9 are important tools for molecular genetics and gene therapy. More broadly, our findings demonstrate the importance of studying how natural and engineered nucleases engage different repair pathways. This work also serves as a potential guide for future engineering efforts by establishing a design paradigm through which genome editors can be biased toward particular repair outcomes.

## Materials and Methods

### Mammalian cell culture

All mammalian cell cultures were maintained in a 37°C incubator at 5% CO_2_. HEK293T human embryonic kidney, HeLa human cervical cancer, A549 human lung cancer, Panc1 human pancreatic cancer, and primary human dermal fibroblast from neonate (GlobalStem) cells were maintained in Dulbecco’s Modified Eagle’s Medium with high glucose, sodium pyruvate, and GlutaMAX (DMEM; ThermoFisher, 10569) supplemented with 10% Fetal Bovine Serum (FBS; ThermoFisher, 10438), and 100 U/mL Penicillin-Streptomycin (ThermoFisher, 15140). Primary human dermal fibroblasts were induced to a quiescent state in medium containing 0.1% Fetal Bovine Serum for 7 days (*35*). For inhibitor studies, cell media was supplemented with 20 µM Rucaparib (MilliporeSigma, PZ0036) or NU7026 (MilliporeSigma, N1537) dissolved in DMSO (MilliporeSigma, D8418).

### Mutagenesis and cloning

Wild-type Cas9 was obtained from pSpCas9 (pX165) and a cloning backbone for gRNA expression was obtained from pX330-U6-Chimeric_BB-CBh-hSpCas9 (pX330), which were gifts from Feng Zhang. Cas9 mutagenesis was performed using PCR-driven splicing by overlap extension using primers listed in Supplementary Table 1. Briefly, one fragment was amplified by PCR from pX165 using the cas9-mut-FWD or cas9-mid-FWD and *mutant*-BOT primers and a second fragment was amplified using the *mutant*-TOP and cas9-mid-REV or cas9-mut-REV primers for each mutant. Each pair of fragments was then spliced by overlap extension PCR using the cas9-mut-FWD and cas9-mid-REV or cas9-mid-FWD and cas9-mut-REV primers to create a Cas9 gene fragment with a single residue mutation. These Cas9 gene fragments were then each cloned back into pX165 using unique BshTI, ApaI, and EcoRI restriction sites to replace the wild-type sequence with the mutant sequence. Additional mutants (double-, triple-, and quadruple-mutants) were made iteratively starting from these single-mutant plasmids. A custom gRNA cloning backbone vector was created by PCR amplification from pX330 using the gRNA-scaffold-NheI-FWD and gRNA-scaffold-EcoRI-REV primers and restriction cloning into pUC19 (ThermoFisher) using NheI and EcoRI digestion. The gRNA spacer sequence oligos, listed in Supplementary Table 2, were phosphorylated with T4 polynucleotide kinase (NEB) and cloned into gRNA cloning backbone by Golden Gate cloning with BpiI digestion. High-fidelity Cas9 variants were obtained from pX165-Cas9-HF1, pX165-eSpCas9, and pX165-HypaCas9, which were gifts from Feng Zhang. Cas9 fusions Cas9-CtIP and Cas9-dn53bp1 were created by restriction cloning of custom geneblocks synthesized by IDT into pX165 at unique KflI and EcoRI restriction sites. Primers were synthesized by IDT. Restriction enzymes were obtained from ThermoFisher. T7 DNA ligase was obtained from NEB. Plasmids were transformed into competent Stbl3 chemically competent *E. Coli* (ThermoFisher). Sequences for the wild-type Cas9, vCas9, and gRNA cloning backbone vectors are presented in the Sequences section.

### Structure analysis

Crystal structures of Cas9 with substrate DNA bound (5F9R) or without substrate DNA bound (4ZT0) were analyzed using PyMol (Schrödinger).

### Cell transfection

Cells were seeded in the maintenance medium without Pen-Strep into 24-well plates at 100,000 cells/well or 48-well plates at 50,000 cells/well. Transfections of HEK293T without repair templates were carried out 24 hrs after seeding using 400 ng Cas9 expression vector and 144 ng gRNA expression vector formulated with 1.36 µL Lipofectamine 2000 (ThermoFisher) at a total volume of 54.4 µL in OptiMEM I (ThermoFisher) per well for 24-well plates, or half these volumes for 48-well plates. Transfections of HEK293T, HeLa, A549, and Panc1 with HDR templates were carried out 24 hrs after seeding using 400 ng Cas9 expression vector, 144 ng gRNA expression vector, and 400 ng ssODN HDR template formulated with 2.11 µL Lipofectamine 2000 at a total volume of 84.4 µL in OptiMEM I per well for 24-well plates, or half these volumes for 48-well plates. Transfections of HEK293T with dual gRNAs were carried out 24 hrs after seeding using 400 ng Cas9 expression vector and 144 ng of each gRNA expression vector formulated with 1.72 µL Lipofectamine 2000 (ThermoFisher) at a total volume of 68.8 µL in OptiMEM I (ThermoFisher) per well for 24-well plates, or half these volumes for 48-well plates. Transfections of HEK293T and primary human dermal fibroblasts with MDR vectors were carried out 24 hrs after seeding (HEK293T) or 7 days after quiescence induction (primary human dermal fibroblasts) using 400 ng Cas9 expression vector, 144 ng gRNA expression vector, and 153-295 ng (equimolar) MDR template formulated with 1.74-2.10 µL (equal volume / DNA) Lipofectamine 2000 at a total volume of 69.7-83.9 µL (equal DNA concentration) in OptiMEM I (ThermoFisher) per well. For sequencing assays, genomic DNA was extracted 72 hrs after transfection using QuickExtract (Epicentre). For flow cytometry assays, cells were transferred to 6-well plates 72 hrs after transfection, split 7 days after transfection, and harvested 10 days after transfection in PBS with 5% FBS (ThermoFisher). Repair templates, listed in Supplementary Table 3, were synthesized by IDT.

### High-throughput sequencing

The targeted loci were amplified from extracted genomic DNA by PCR using Herculase II polymerase (Agilent). The PCR primers included Illumina sequencing handles as well as replicate-specific barcodes. These PCR products were then tagged with sample-specific barcodes and sequenced on an Illumina MiSeq. Primers, listed in Supplementary Table 4, were synthesized by IDT.

#### Sanger sequencing

The targeted loci were amplified from extracted genomic DNA by PCR using Herculase II polymerase (Agilent). PCR amplicons were sequenced using primers ∼200 bp from the expected cut site. To measure editing frequencies, the sequencing traces were analyzed using TIDE (*36*). Primers, listed in Supplementary Table 5, were synthesized by IDT.

#### Cell cycle profiling

Primary human dermal fibroblasts were seeded in the maintenance medium without Pen-Strep into 12-well plates at 100,000 cells/well (dividing cells) or 200,000 cells/well (quiescent cells). Profiling was carried out 24 hrs after seeding (dividing cells) or 7 days after quiescence induction (quiescent cells). Cells were pulsed with 10 µM 5-ethynyl-2’-deoxyuridine (EdU) added to the medium for 1 hr before harvesting. Cells from four wells per condition were pooled, fixed, permeabilized, and stained for EdU incorporation using a Click-iT Plus EdU Alexa Fluor 647 Flow Cytometry Assay Kit (ThermoFisher). To label total DNA content, cells were resuspended in Click-iT saponin-based permeabilization and wash buffer (ThermoFisher) with 20 µg/mL propidium iodide and 100 µg/mL RNase A for 30 min. Cells were analyzed by flow cytometry to profile cell cycle stage.

#### Flow cytometry

Flow cytometry analysis was performed on an LSR Fortessa analyzer and data was collected using FACSDiva (BD Biosciences). Cells were first gated comparing SSC-A and FSC-A, then SSC-H and SSC-W, then FSC-H and FSC-W parameters to select for single cells. To assess editing frequencies, cells were gated for GFP (488 nm laser excitation, 530/30 nm filter detection) and BFP (405 nm laser excitation, 450/50 nm filter detection). To profile cell cycle stage, cells were gated for propidium iodide (561 nm laser excitation, 610/20 nm filter detection) and Alexa Fluor 647 (640 nm laser excitation, 670/30 nm filter detection). Flow cytometry data were analyzed using FlowJo (FlowJo).

### Genome editing analysis

To measure editing outcomes, the high-throughput sequencing data were analyzed using CRISPResso2 (*37*). Total editing rates were quantified as the fraction of edited reads out of total sequencing reads. Indel rates were quantified as the fraction of reads containing indels out of total sequencing reads. Precise editing rates were quantified as the fraction of reads containing a perfect match to the expected edit out of total sequencing reads. Frequencies of specific indel sizes were quantified as the fraction of reads containing these sizes out of all edited reads. Depletion of specific indel sizes by templated repair was quantified as the fractional reduction in the frequency of that indel size, comparing frequencies for when a template was present versus absent. Mean indel sizes were calculated as the mean of the absolute values of indel sizes weighted by their indel fractions.

#### DNA break structure analysis

To measure DNA break structures for Cas9 variants, editing outcomes for dual-gRNA cutting of genomic DNA were analyzed as previously described (*31*). HEK293T cells were edited with pairs of gRNAs targeting the *EMX1* (EMX1 gRNA 1 and gRNA 2) or *CXCR4* (CXCR4 gRNA 1 and gRNA 2) loci. The gRNA pairs were complementary to the same strand at each locus and were expected to make cuts 84bp apart, resulting in large precise deletions. The loci were amplified and sequenced by high-throughput sequencing. The high-throughput sequencing data were analyzed using CRISPResso2 (*37*), using the expected 84bp deletion junction as a reference sequence. To assess DNA break structure, sequencing reads aligned to the deletion junction reference were analyzed for insertion sequences perfectly matching the sequences flanking the expected gRNA cut sites. The positions of these matching sequences at the two gRNA sites were used to determine cut positions leading to each read. Frequencies of these cut positions were quantified as the fraction of reads resulting from these specific cut positions out of all reads containing the deletion junctions with or without insertions.

#### Repair pathway outcome analysis

For repair pathway analysis, next-generation sequencing reads were classified using annotated repair were analyzed using CRISPResso2 (*37*). The same gRNA and locus sequences were also analyzed using Indelphi to identify whether each predicted indel was associated with a microhomology (MMEJ) or not (NHEJ), along with microhomology sizes. These repair pathway labels for each edited sequence from Indelphi analysis were then applied to the matching sequencing reads for the editing experiment. Frequencies of NHEJ, MMEJ, and precise editing were quantified as the fraction of reads containing these types of edits out of all edited reads.

#### Off-target activity analysis

To assess off-target cutting activity, indel rates were analyzed at known off-target sites previously reported for two gRNAs (EMX1 gRNA 2 and VEGFA gRNA 1) (*24*). Indel rates were determined by analysis of Sanger sequencing traces at these on-target and off-target loci using TIDE (*36*).

#### Statistical analysis

Specific statistical comparisons are indicated in the figure legends. Error bars indicate the standard error for three independent replicates. In most comparisons, significance was assessed using unpaired, two-tailed Student’s t-tests. For correlations, significance was assessed using Pearson’s tests. For linear regressions, significance was assessed using ANCOVA tests.

## Acknowledgements

The authors thank Salil Garg, Dig Bijay Mahat, Luke Rhym, and Amanda Whipple for helpful discussions.

## Author Contributions

V.P.C., P.A.S., and R.L. conceived or designed the work; V.P.C., P.A.S., and R.L. acquired, analyzed, or interpreted the data; V.P.C., P.A.S., and R.L. drafted the work or substantively revised it.

## Data and Materials Availability

Raw next-generation sequencing data are to be made available on NCBI SRA. Plasmids are to be deposited on Addgene.

## Supplementary Materials

### Supplementary Figures

**Figure S1.**
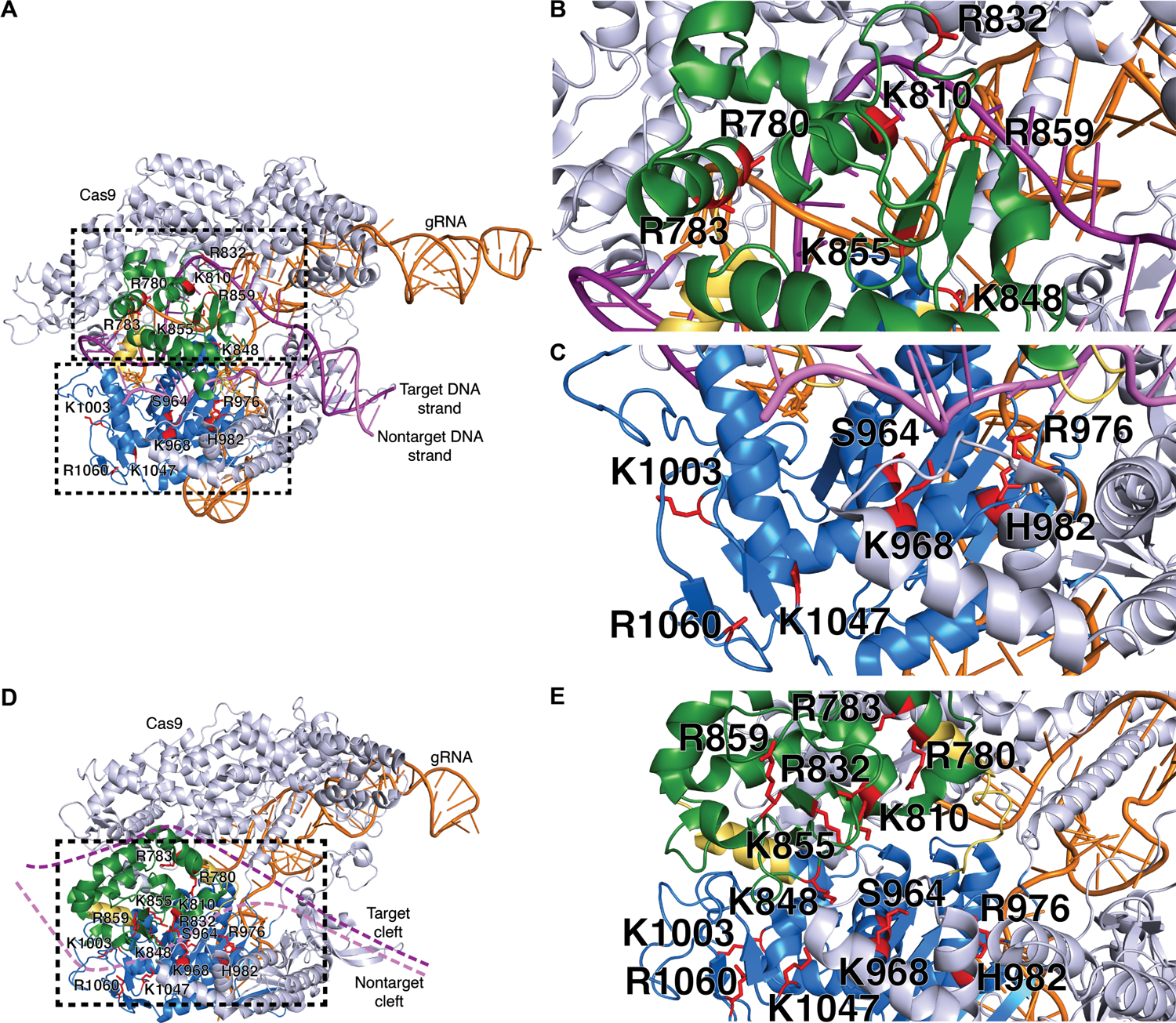
Detailed views of mutated Cas9 residues. (**A**) Wide view of Cas9 residues at the interface with the substrate DNA strands that were selected for mutation and screening. The mutated residues (red) are located in either the mobile HNH domain (green) or the immobile RuvC domain (blue), which are connected by linkers (yellow). (**B**) Focused view of the residues from **A** in the target cleft. (**C**) Focused view of the residues from **A** in the nontarget cleft. (**D**) Location of the same Cas9 residues in a structure of Cas9 without the DNA substrate bound. (**E**) Focused view of the residues from **D** in the nontarget cleft.

**Figure S2.**
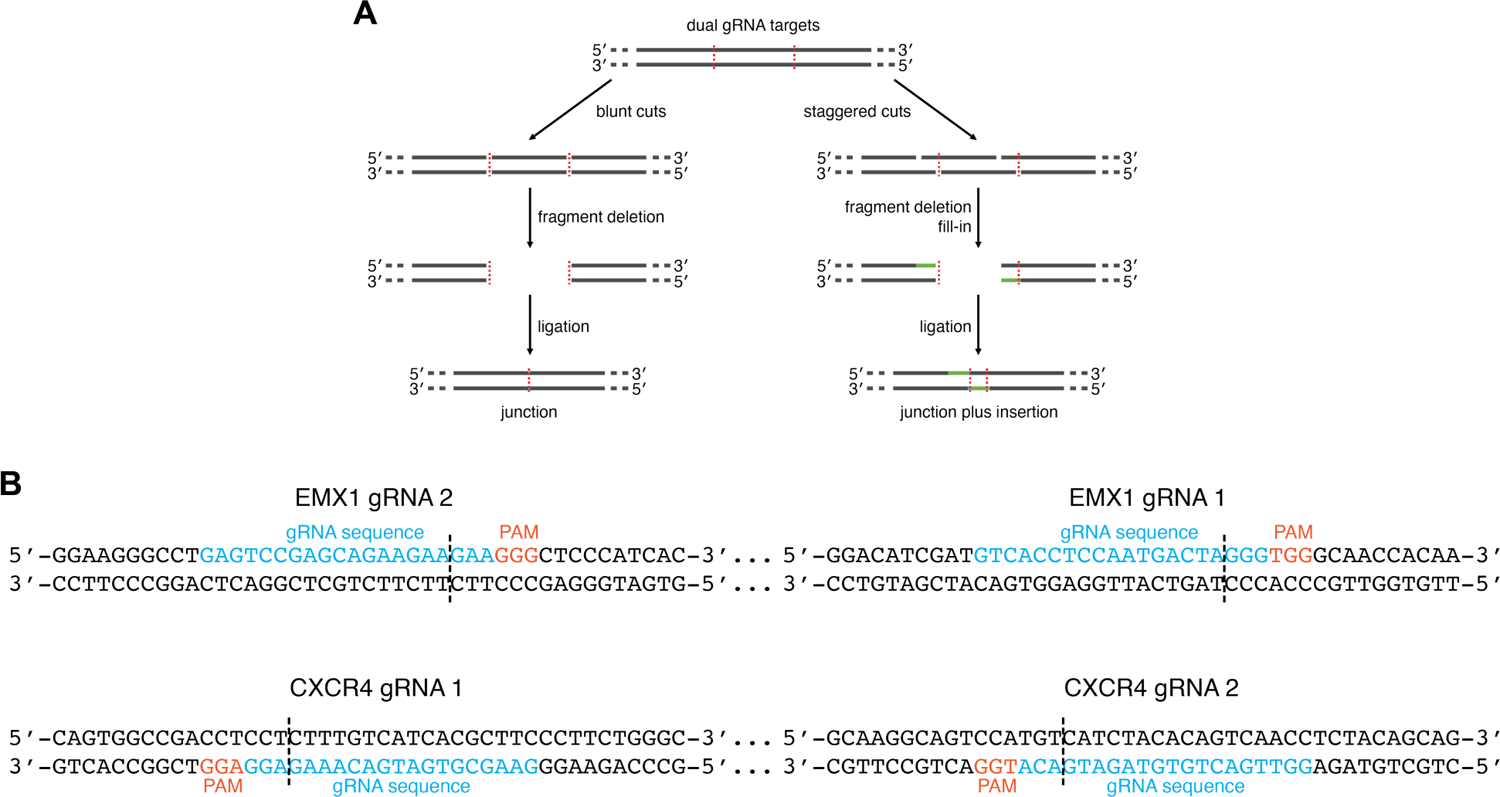
Assay for determining DNA break structure induced by Cas9 variants. (**A**) Assay for DNA break structure, where paired DSBs lead to perfect deletion junctions for blunt cuts and insertions in the deletion junctions for staggered cuts. The sequences of the insertions indicate cut positions in each strand. (**B**) Design of gRNA pairs to introduce deletion junctions.

**Figure S3.**
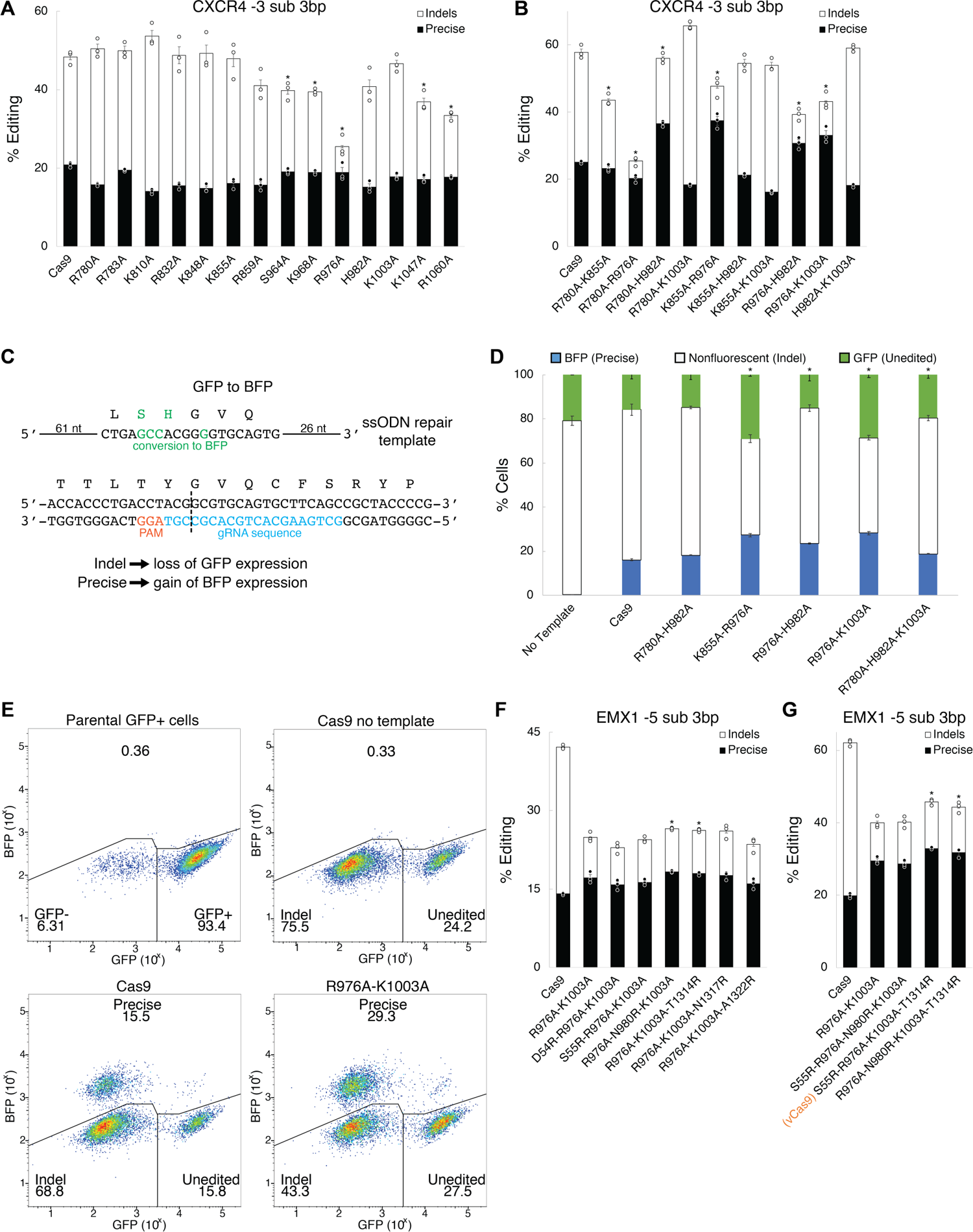
Rational engineering produces Cas9 variants with increased HDR frequency. (**A, B**) Screen of precise editing and indel frequencies for engineered Cas9 (**A**) single-mutant and (**B**) double-mutant variants using an HDR template. (**C**) Design of a gRNA and HDR template to introduce a GFP to BFP conversion. (**D**) Frequency of precise gene conversion from GFP to BFP. (**E**) Flow cytometry plots showing BFP-positive (indicating precise edits), nonfluorescent (indicating indels), and GFP-positive (indicating unedited cells) cell population fractions. (**F**, **G**) Screen of precise editing and indel frequencies for engineered Cas9 (**F**) triple-mutant and (**G**) quadruple-mutant variants using an HDR template. * indicates p < 0.05 for precise editing frequency compared to wild-type Cas9 in **A**, **B**, **D** or for total editing frequency compared to wild-type Cas9 in **F**, **G**. Data were analyzed by deep sequencing in **A**, **B**, **F**, **G** or flow cytometry in **D**, **E** and represent means of *n* = 3 independent replicates with standard errors.

**Figure S4.**
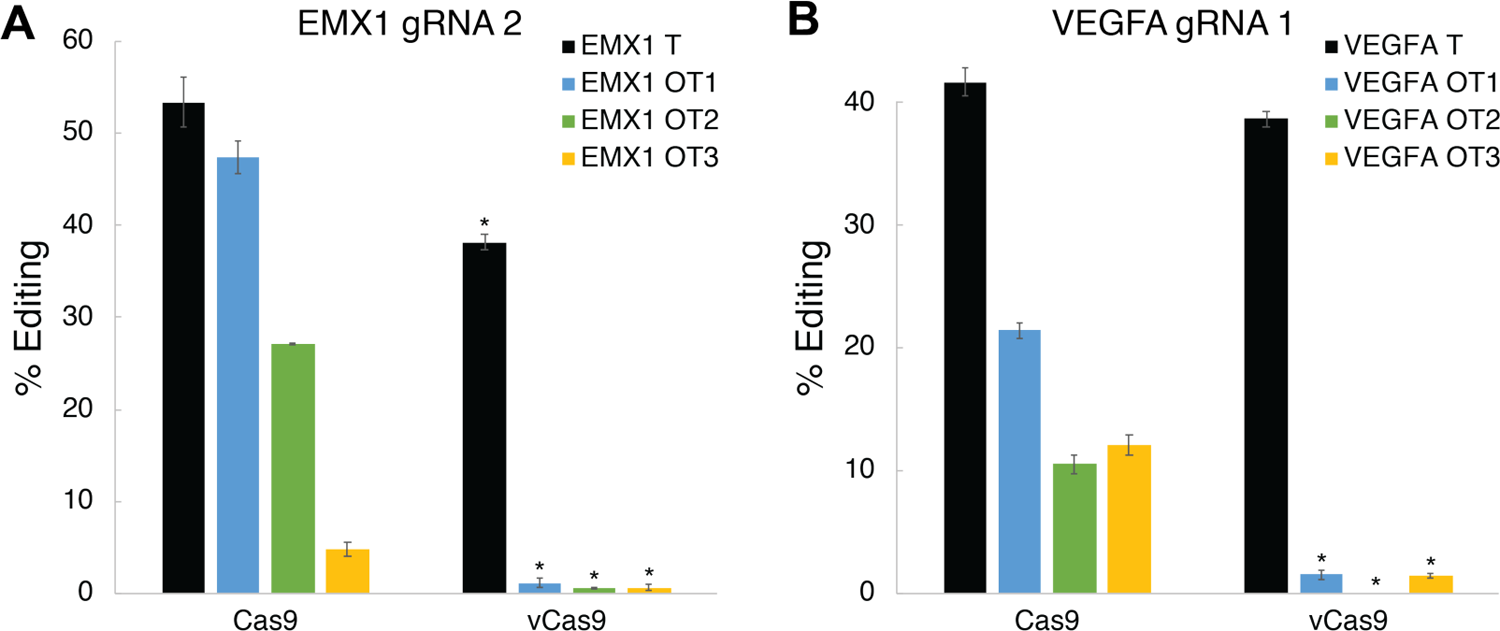
vCas9 improves on-target versus off-target editing specificity. (**A**, **B**) Indel frequencies at on-target (T) and off-target (OT1-OT3) loci. * indicates p < 0.05 for indel frequency compared to wild-type Cas9. Data were analyzed by Sanger sequencing and represent means of *n* = 2-3 independent replicates with standard errors.

**Figure S5.**
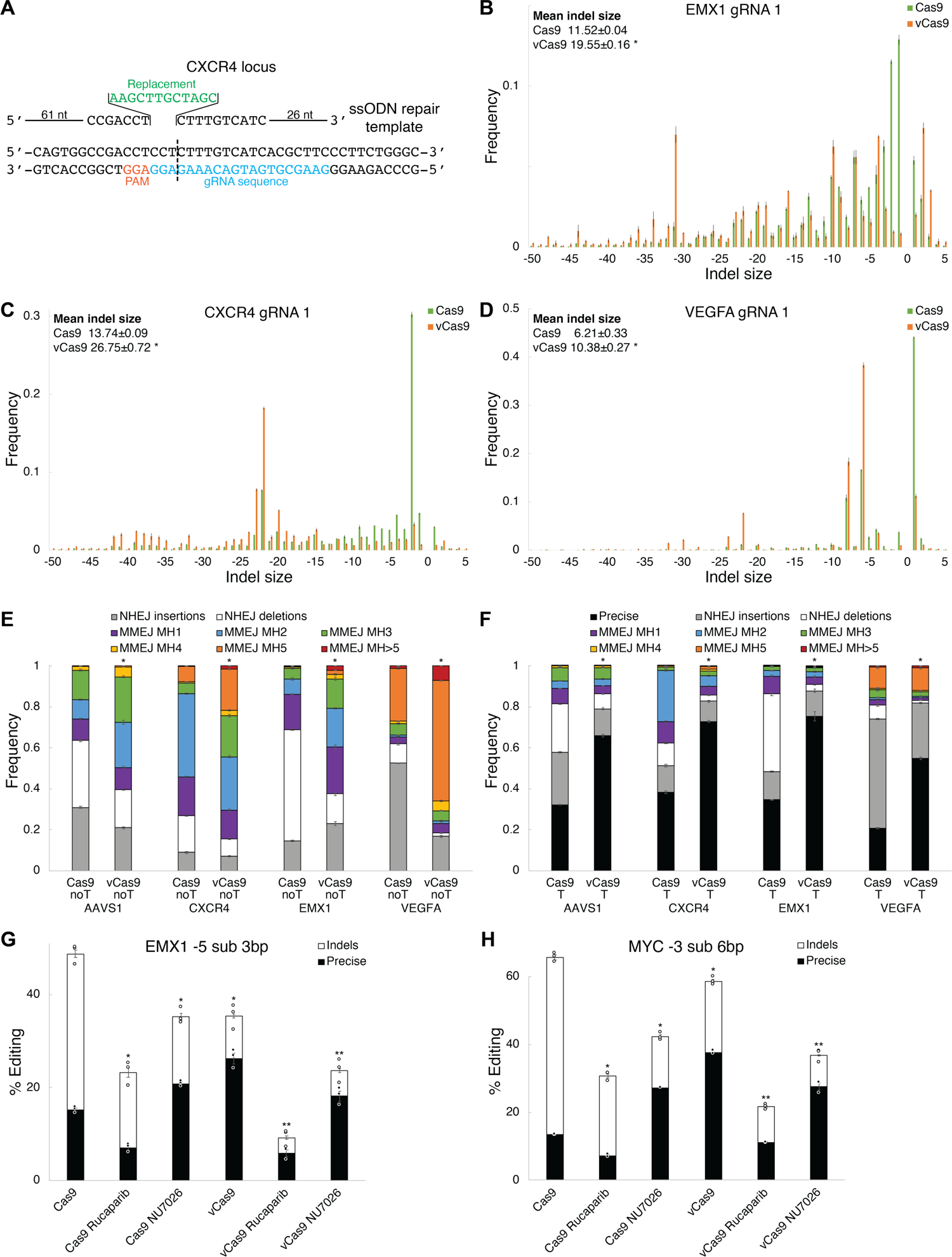
Engineered Cas9 variants suppress small NHEJ indels and promote larger MMEJ indels. **(A)** Design of a gRNA and HDR template to introduce precise edits. (**B**-**D**) Distributions of indel sizes induced. (**E**, **F**) Frequencies of repair pathways engaged (**E**) without (noT) and (**F**) with (T) a repair template at several loci. NHEJ mutations are subset into insertions or deletions, while MMEJ mutations are subset by microhomology length (MH1 to MH>5). (**G**, **H)** Precise editing and indel frequencies using HDR templates without or with an MMEJ inhibitor (Rucaparib) or NHEJ inhibitor (NU7026). * indicates p < 0.05 for indel size compared to wild-type Cas9 in **B**-**D**, for NHEJ frequency compared to wild-type Cas9 in **E**, **F**, or for precise editing frequency compared to wild-type Cas9 in **G**, **H**. ** indicates p < 0.05 for precise editing frequency compared to vCas9. Data were analyzed by deep sequencing and represent means of *n* = 3 independent replicates with standard errors.

**Figure S6.**
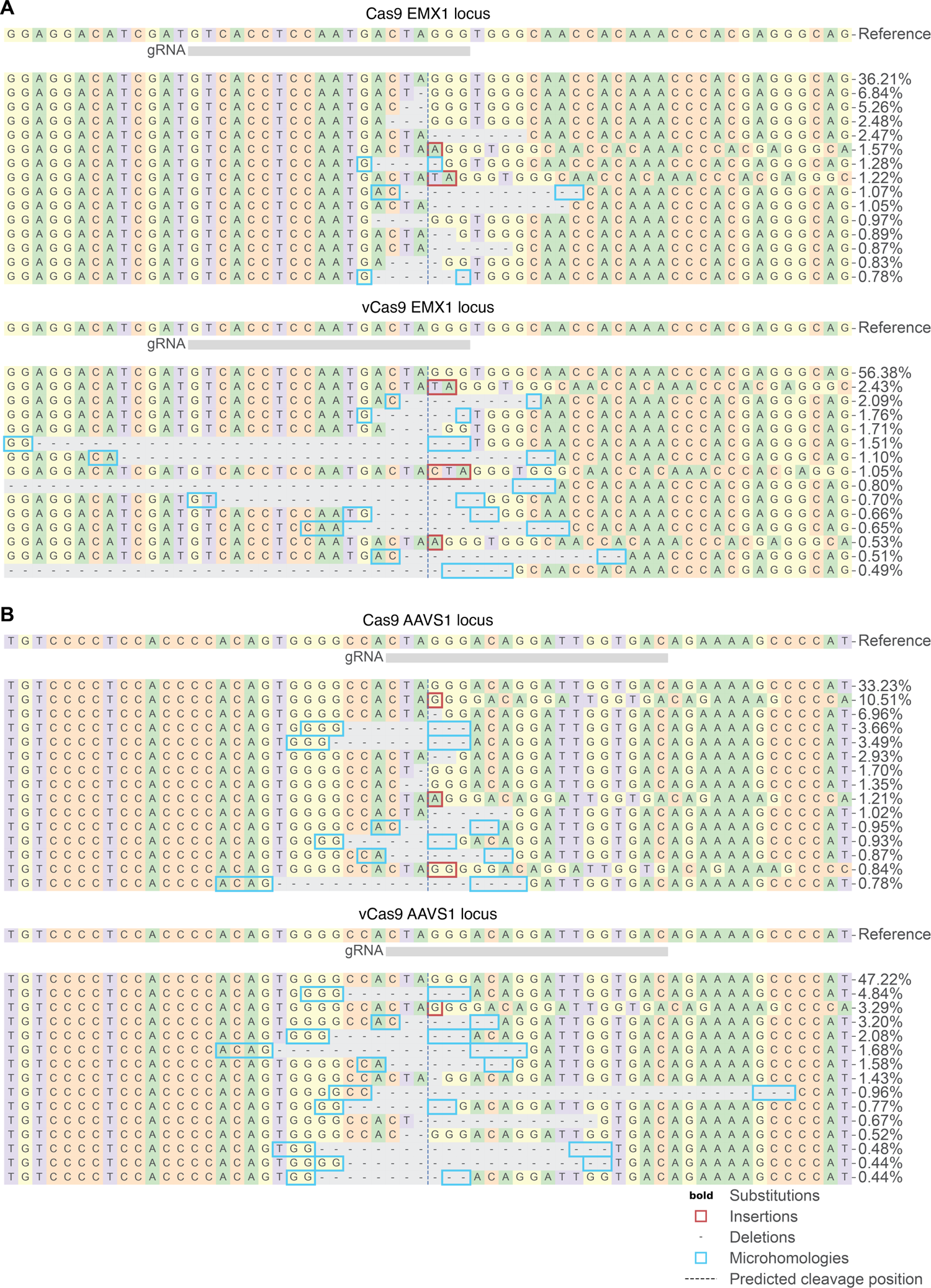
vCas9 suppresses NHEJ and promotes MMEJ. (**A**, **B**) Rates of the top sequences resulting from editing with Cas9 variants at the (**A**) EMX1 and (**B**) AAVS1 loci. Substitutions, insertions, and deletions are depicted. Deletions at microhomologies are labeled. The most frequent fifteen sequences are displayed along with percentages out of all sequencing reads. Data were analyzed by deep sequencing and represent a single replicate.

**Figure S7.**
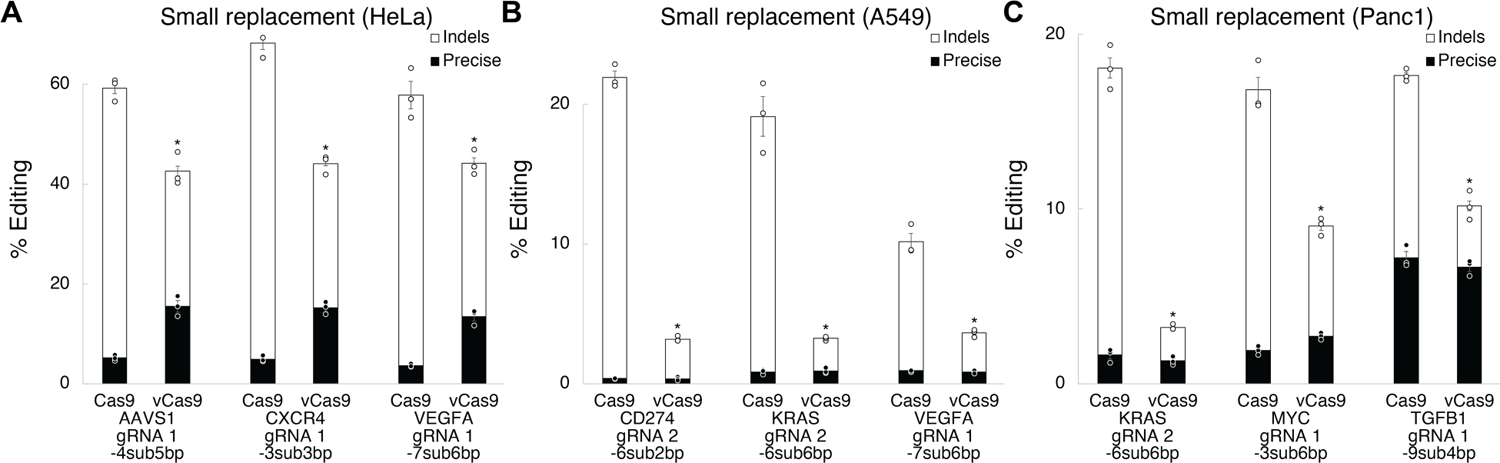
vCas9 enhances precise editing and suppresses indels across cell types. (**A-C**) Precise editing and indel frequencies using small replacement (<10 bp) HDR templates at several loci in (**A**) HeLa, (B) A549, and (**C**) Panc1 cells. * indicates p < 0.05 for precise editing frequency compared to wild-type Cas9. Data were analyzed by deep sequencing and represent means of *n* = 3 independent replicates with standard errors.

**Figure S8.**
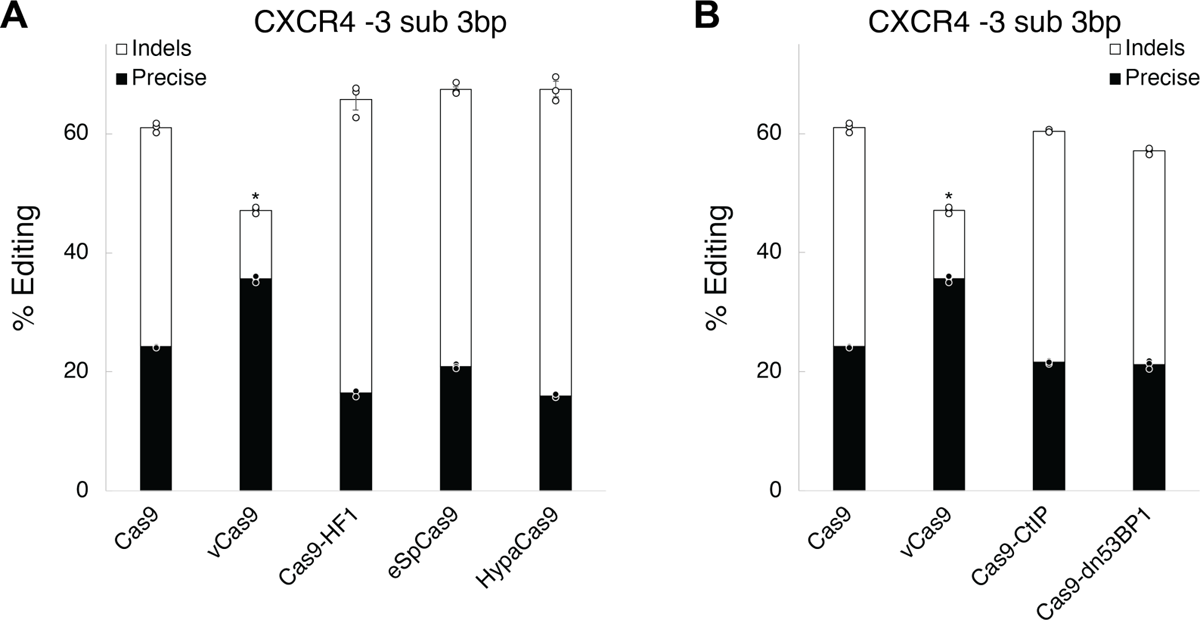
vCas9 enhances precise editing versus other Cas9 variants and fusions. (**A, B**) Precise editing and indel frequencies using an HDR template. * indicates p < 0.05 for precise editing frequency compared to wild-type Cas9. Data were analyzed by deep sequencing and represent means of *n* = 3 independent replicates with standard errors.

**Figure S9.**
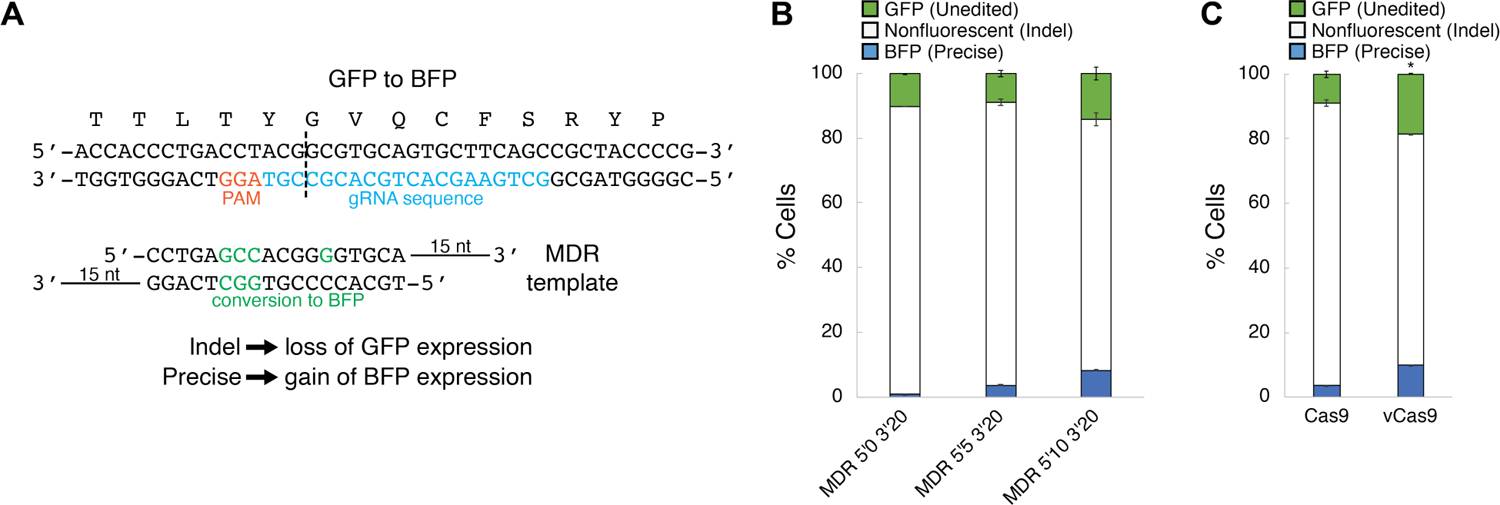
MDR enables efficient precise editing through MMEJ. (**A**) Design of a gRNA and MDR template to introduce a GFP to BFP conversion. (**B**) Precise editing and indel frequencies using MDR templates with varied microhomology arm lengths (0-10 bp at the 5’ ends, 20 bp at the 3’ ends). (**C**) Frequency of precise gene conversion from GFP to BFP. Data were analyzed by flow cytometry and represent means of *n* = 3 independent replicates with standard errors.

**Figure S10.**
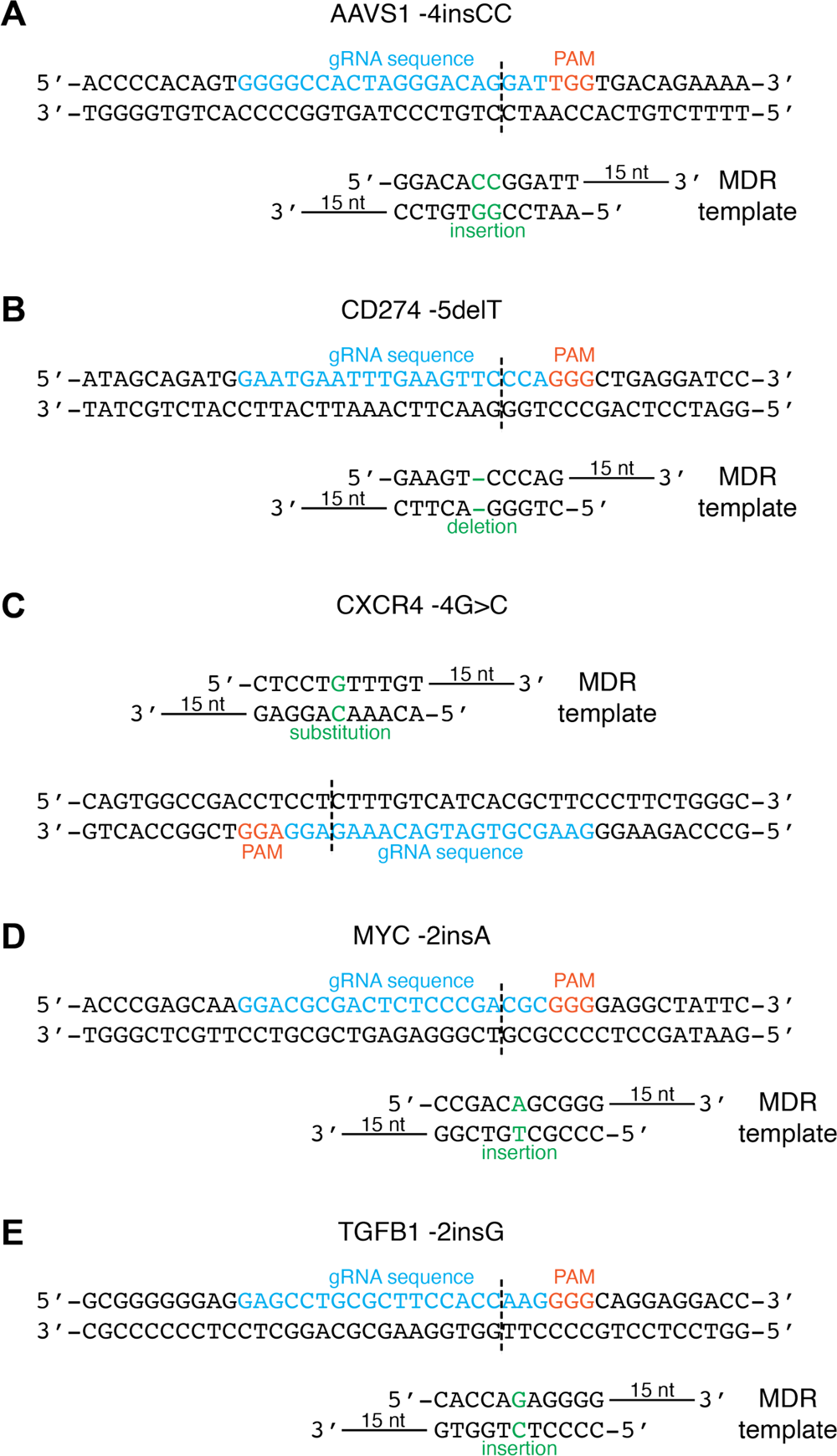
Additional designs for precise editing through MMEJ by MDR. (**A-E**) Design of gRNAs and MDR templates to introduce precise edits.

**Figure S11.**
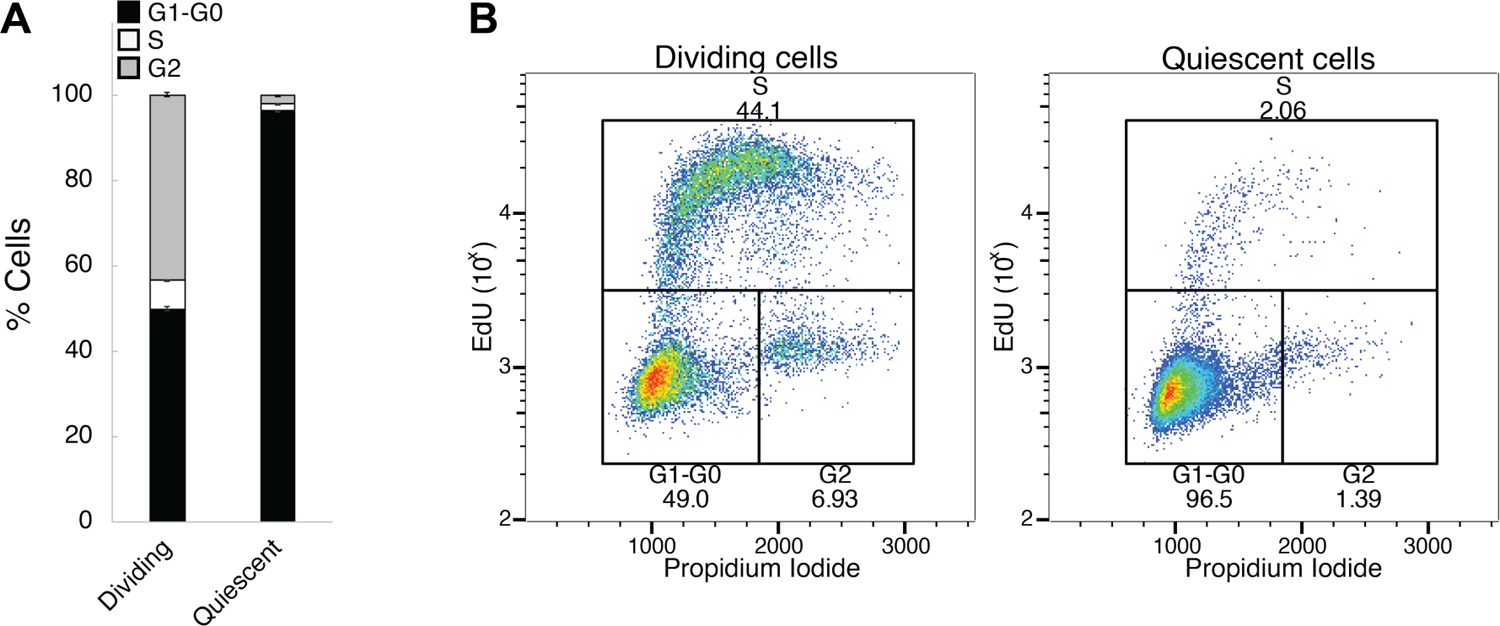
Quiescent primary human dermal fibroblasts are nearly all non-dividing. (**A**) Cell cycle profiling of dividing and quiescent primary human dermal fibroblasts. (**B**) Flow cytometry plots showing EdU-high (indicating S phase), Propidium Iodide-high (indicating G2 phase), and EdU-low Propidium Iodide-low (indicating G1 or G0 phase) cell population fractions. All data were analyzed by flow cytometry and represent means of *n* = 3 independent replicates with standard errors.

### Supplementary Tables

**Supplementary Table 1.**
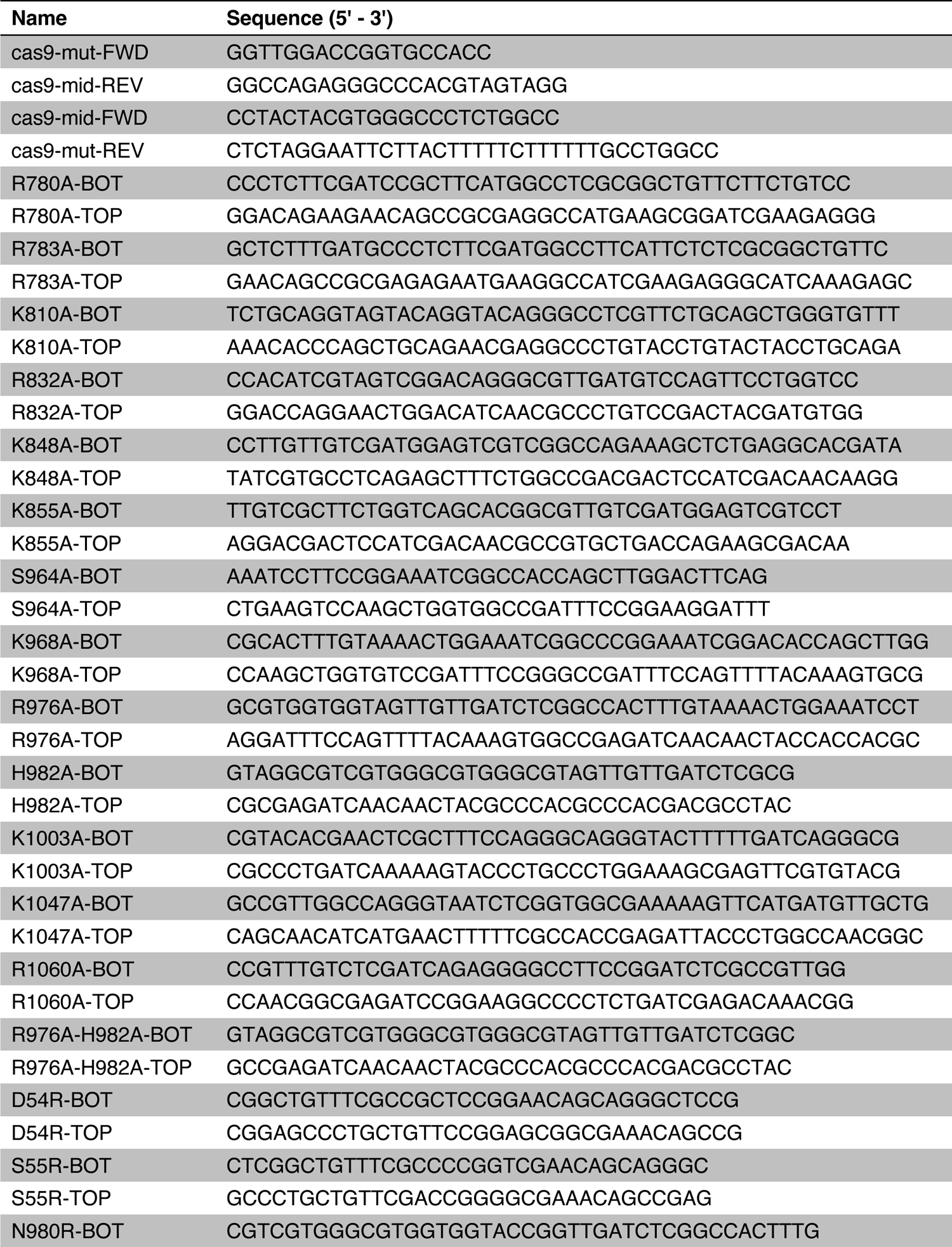
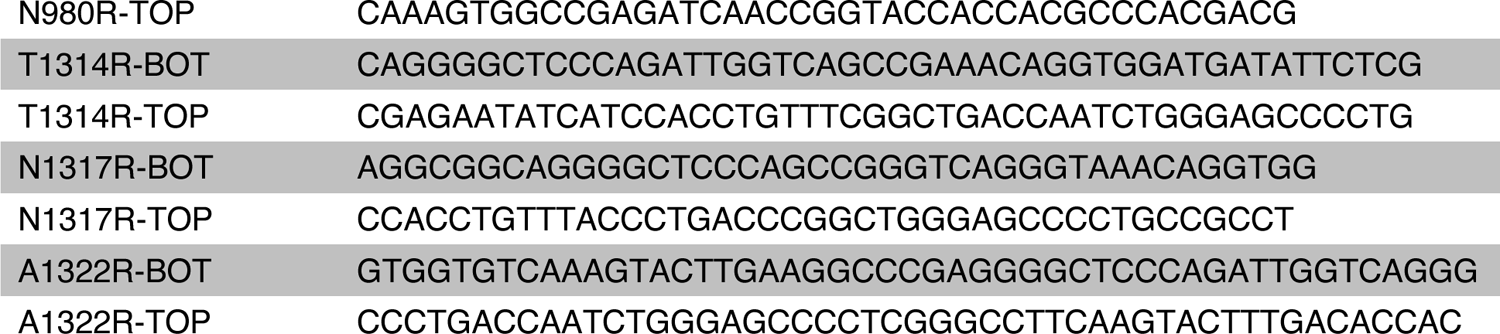
Oligodeoxynucleotide sequences used for mutagenesis and cloning.

**Supplementary Table 2.**
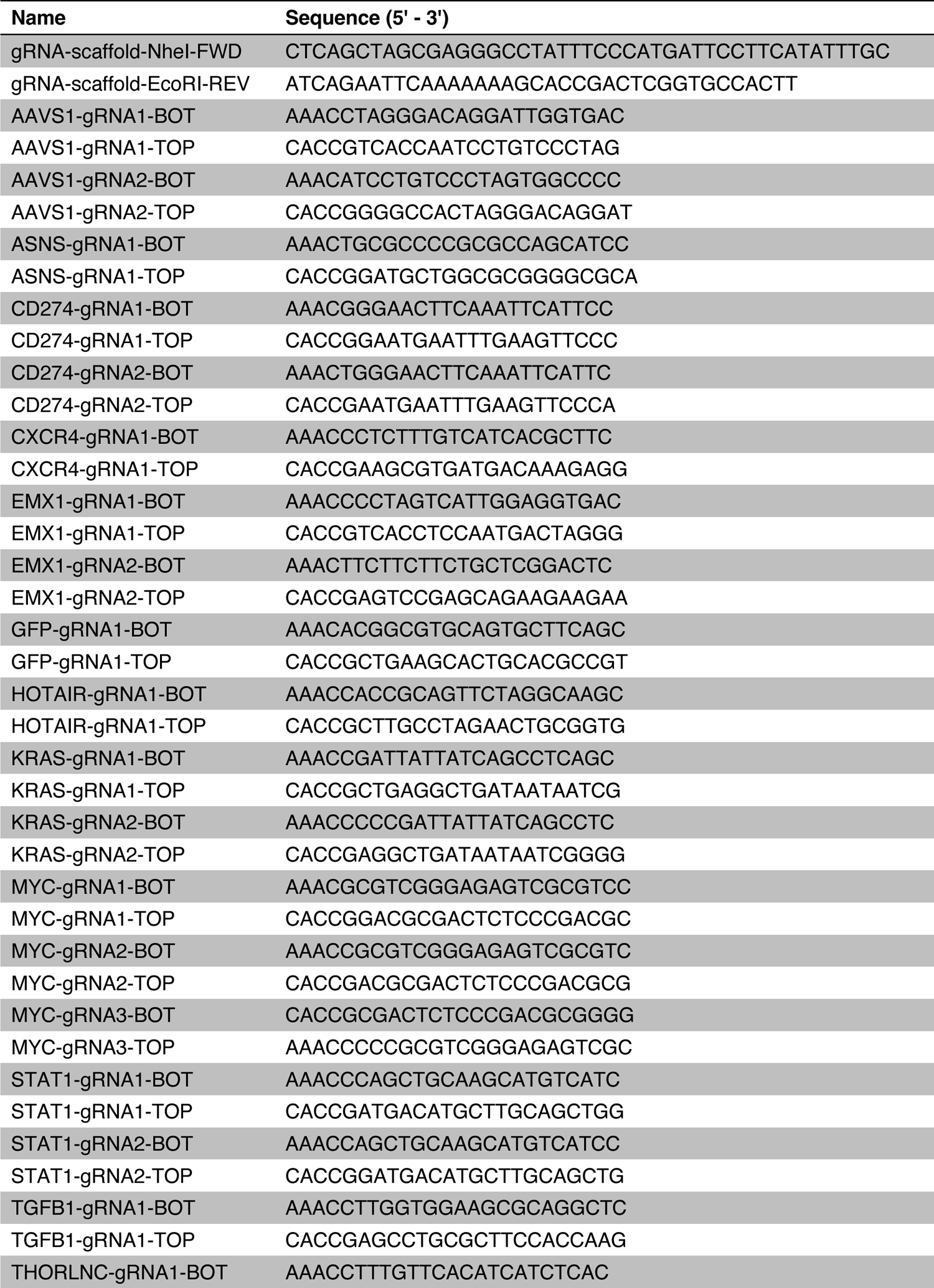
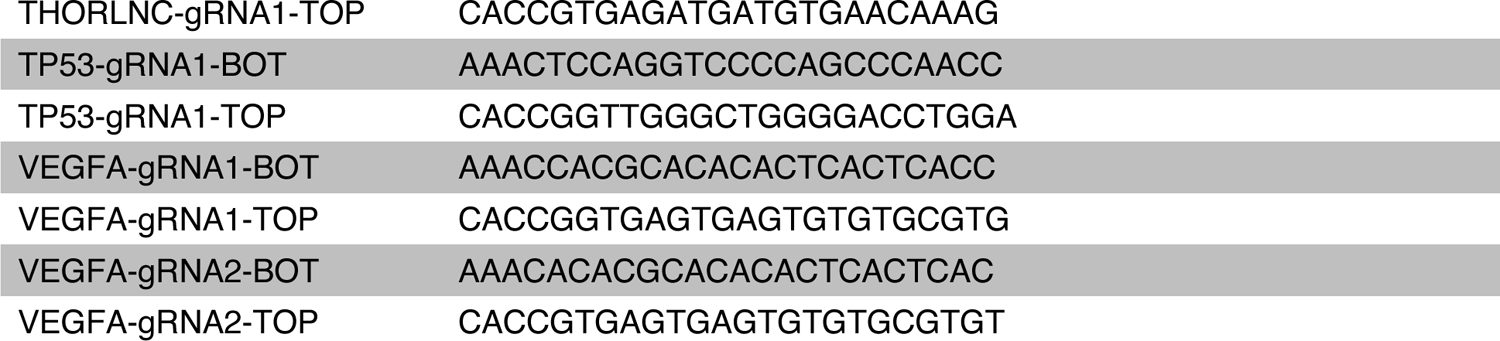
Oligodeoxynucleotide sequences used for gRNA cloning.

**Supplementary Table 3.**
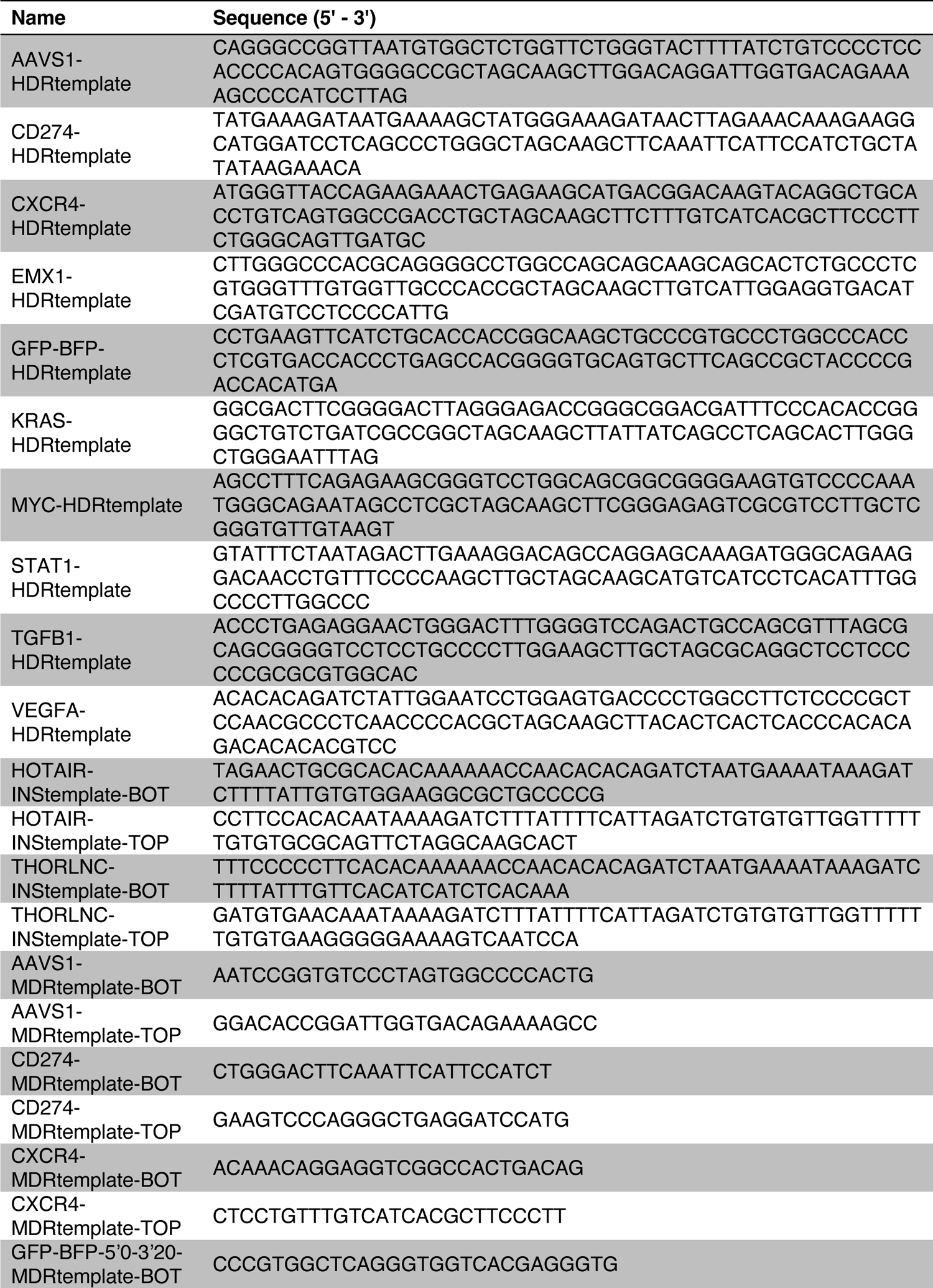
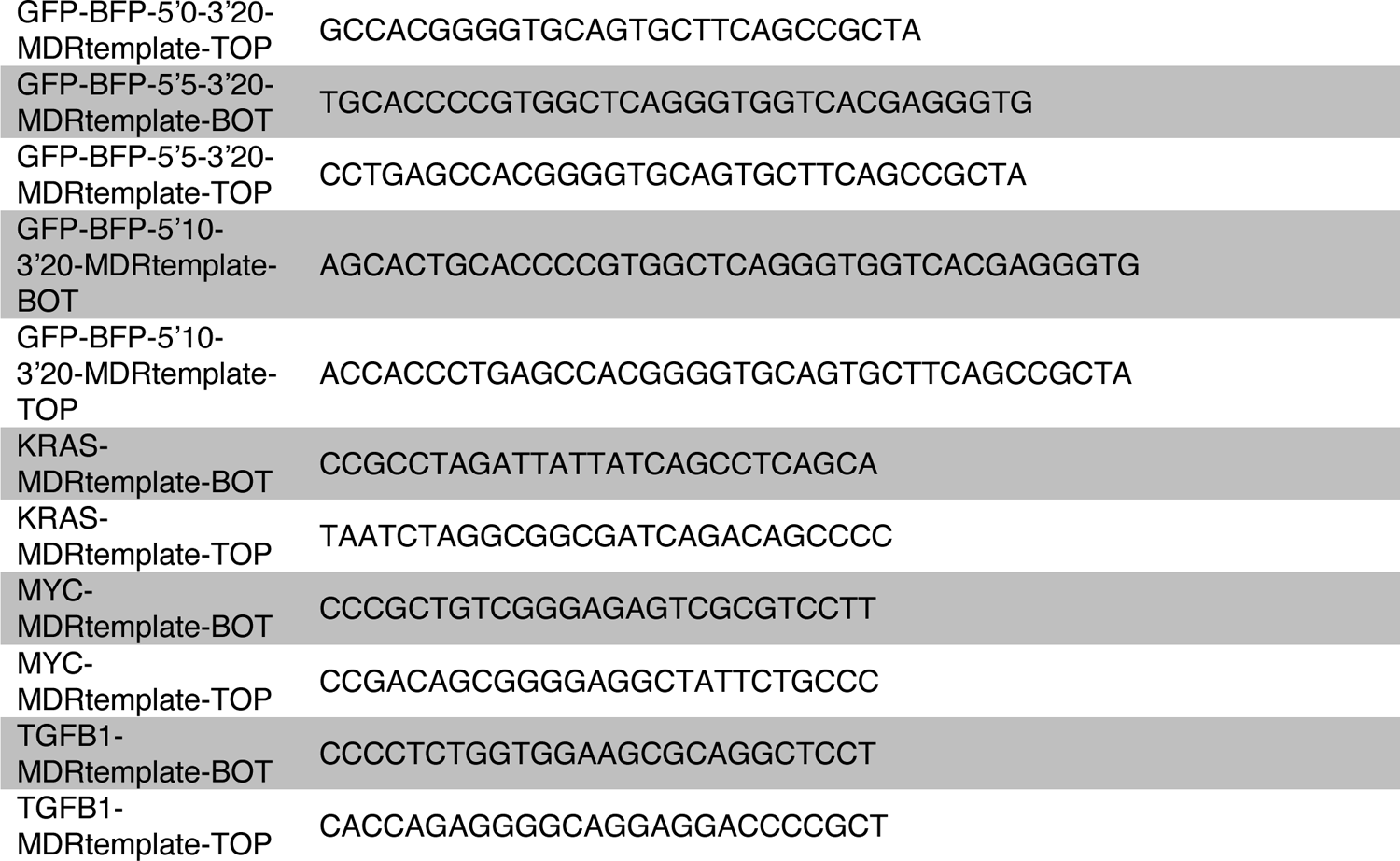
Oligodeoxynucleotide sequences used for HDR and MDR templates.

**Supplementary Table 4.**
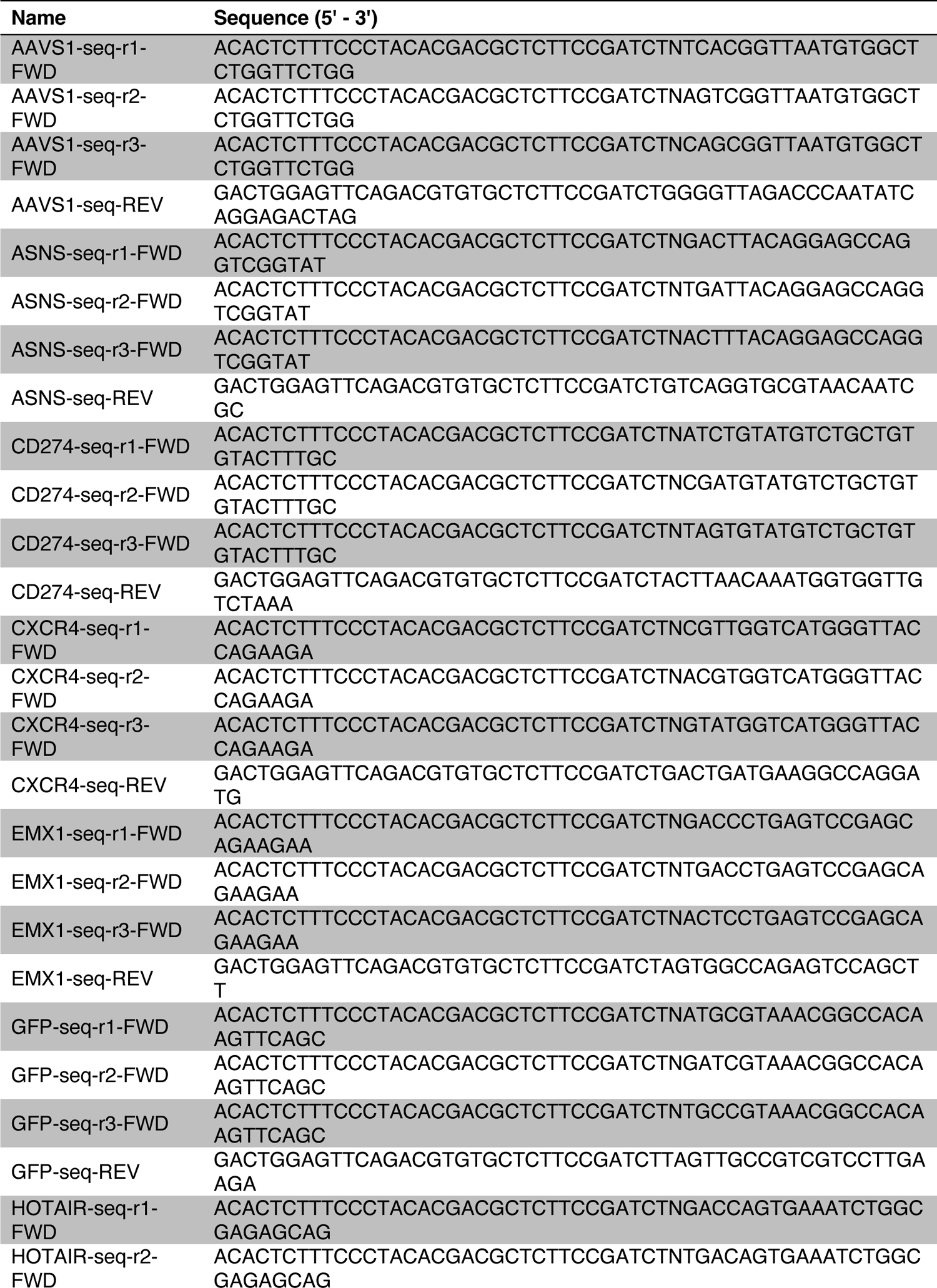
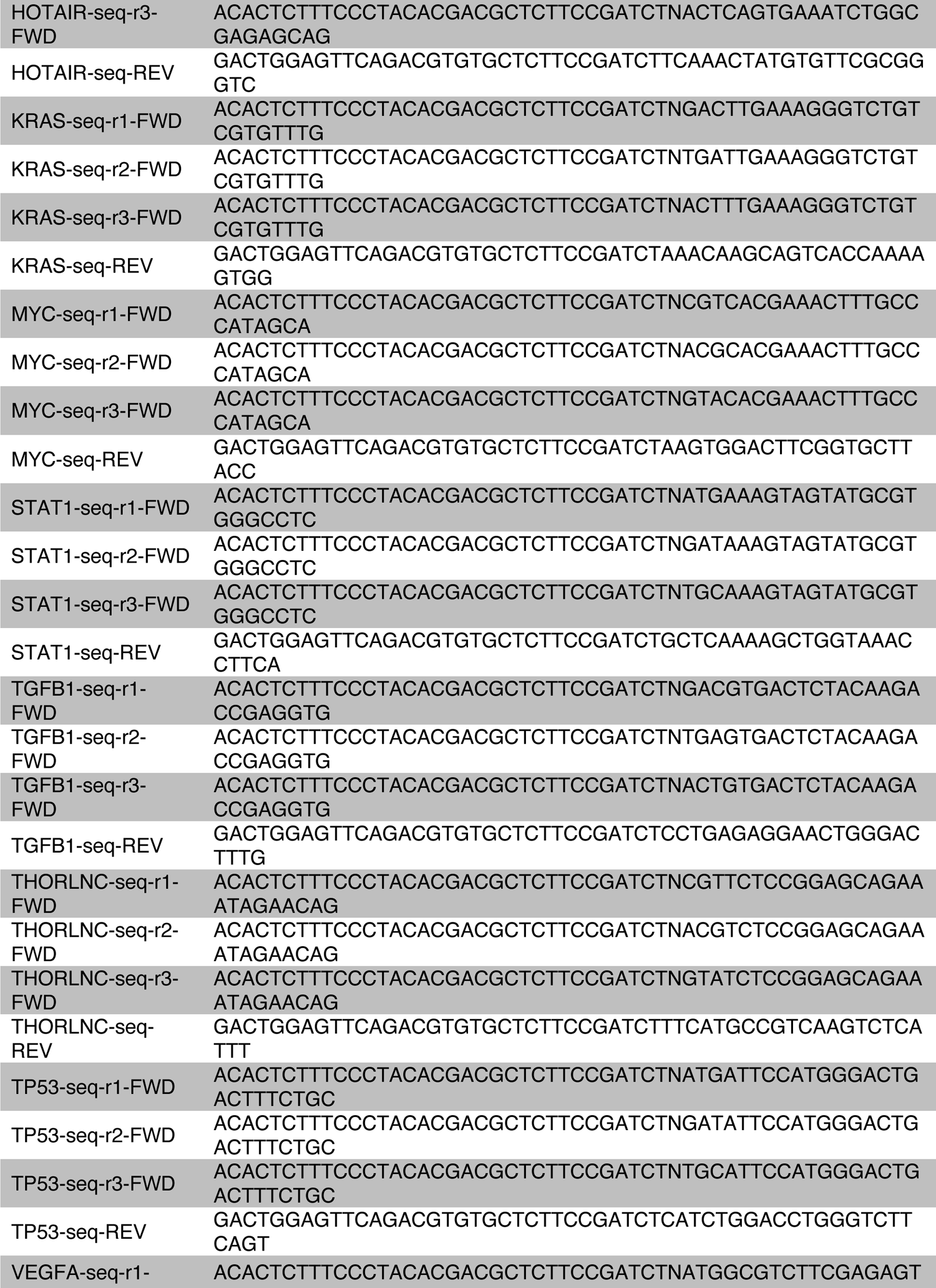
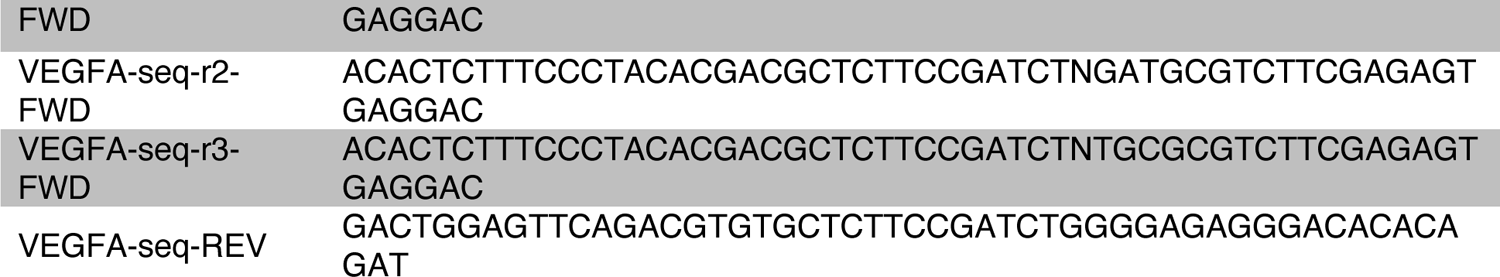
Oligodeoxynucleotide sequences used for next-generation sequencing.

**Supplementary Table 5.**
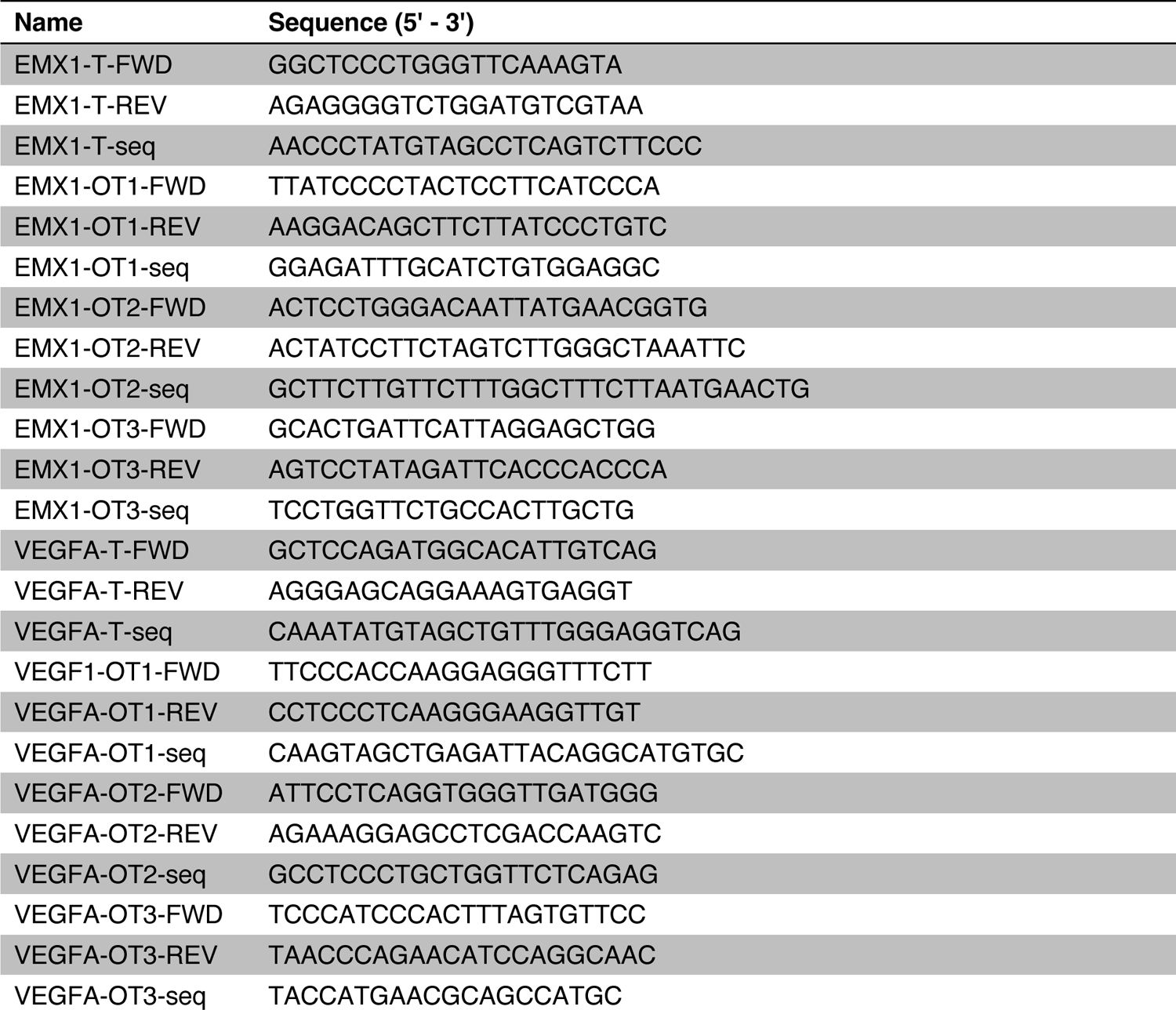
Oligodeoxynucleotide sequences used for Sanger sequencing.

### Sequences

#### Cas9

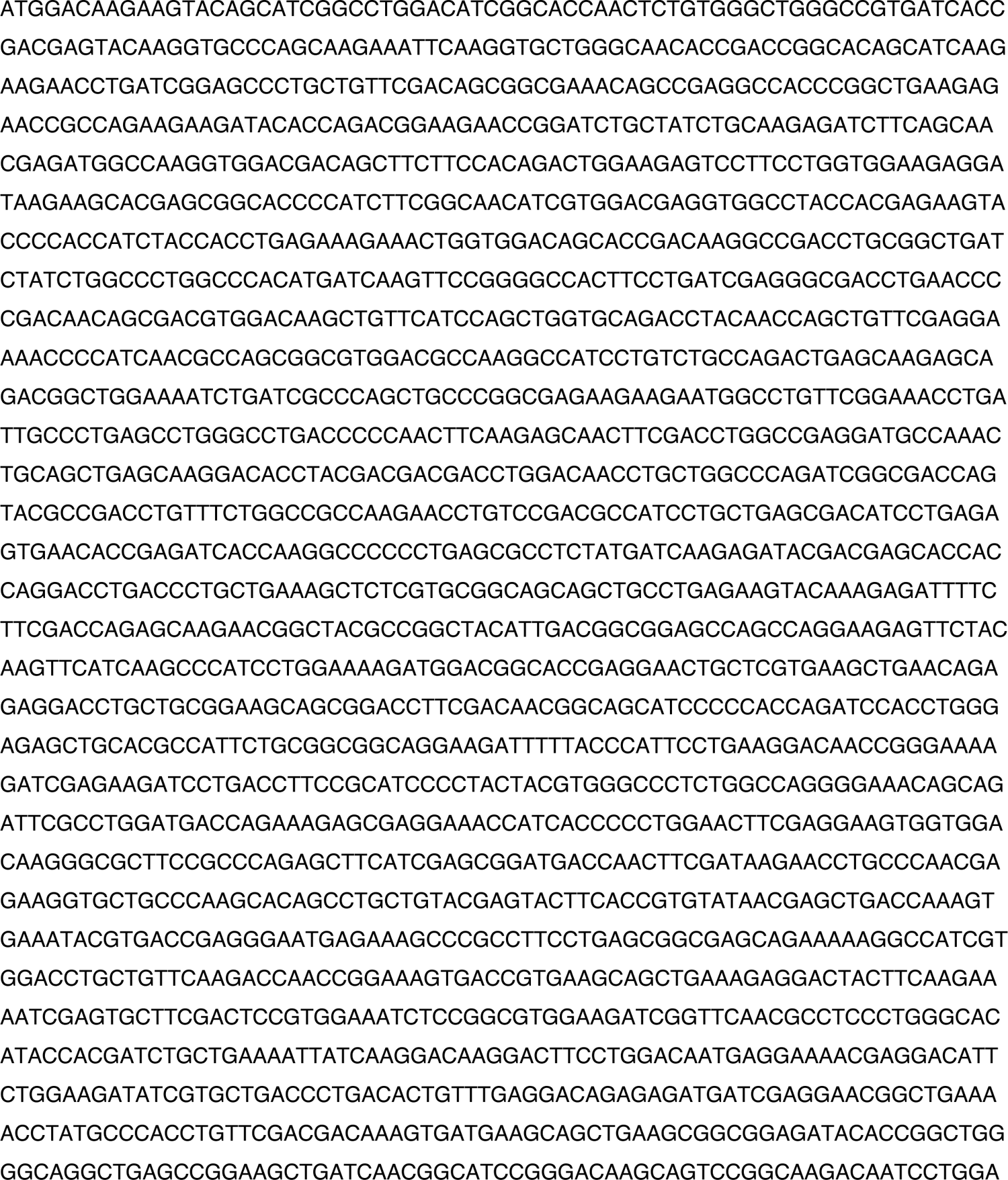
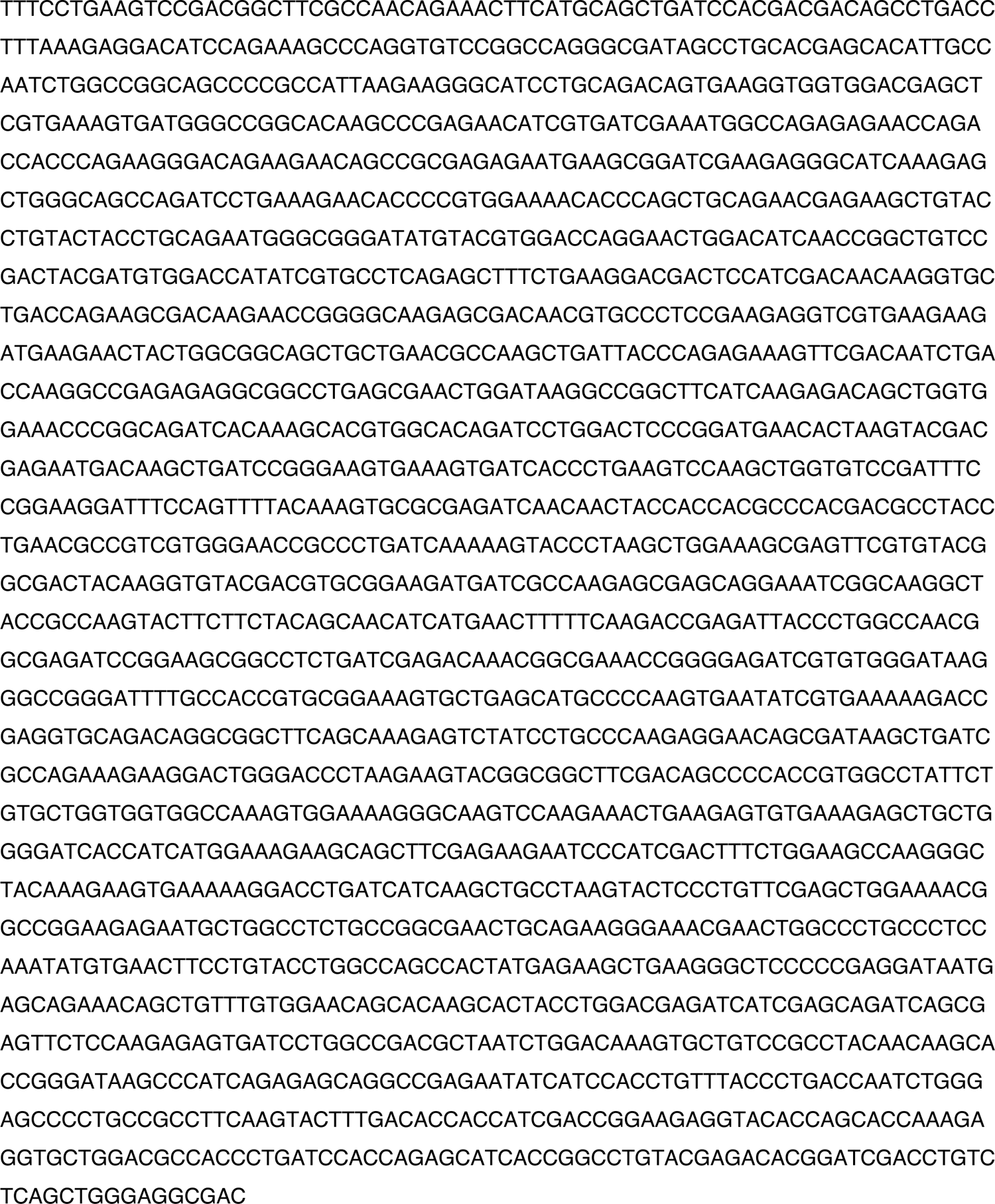

#### Cas9^S55R-R976A-K1003A-T1314R^ (vCas9), with mutations in red

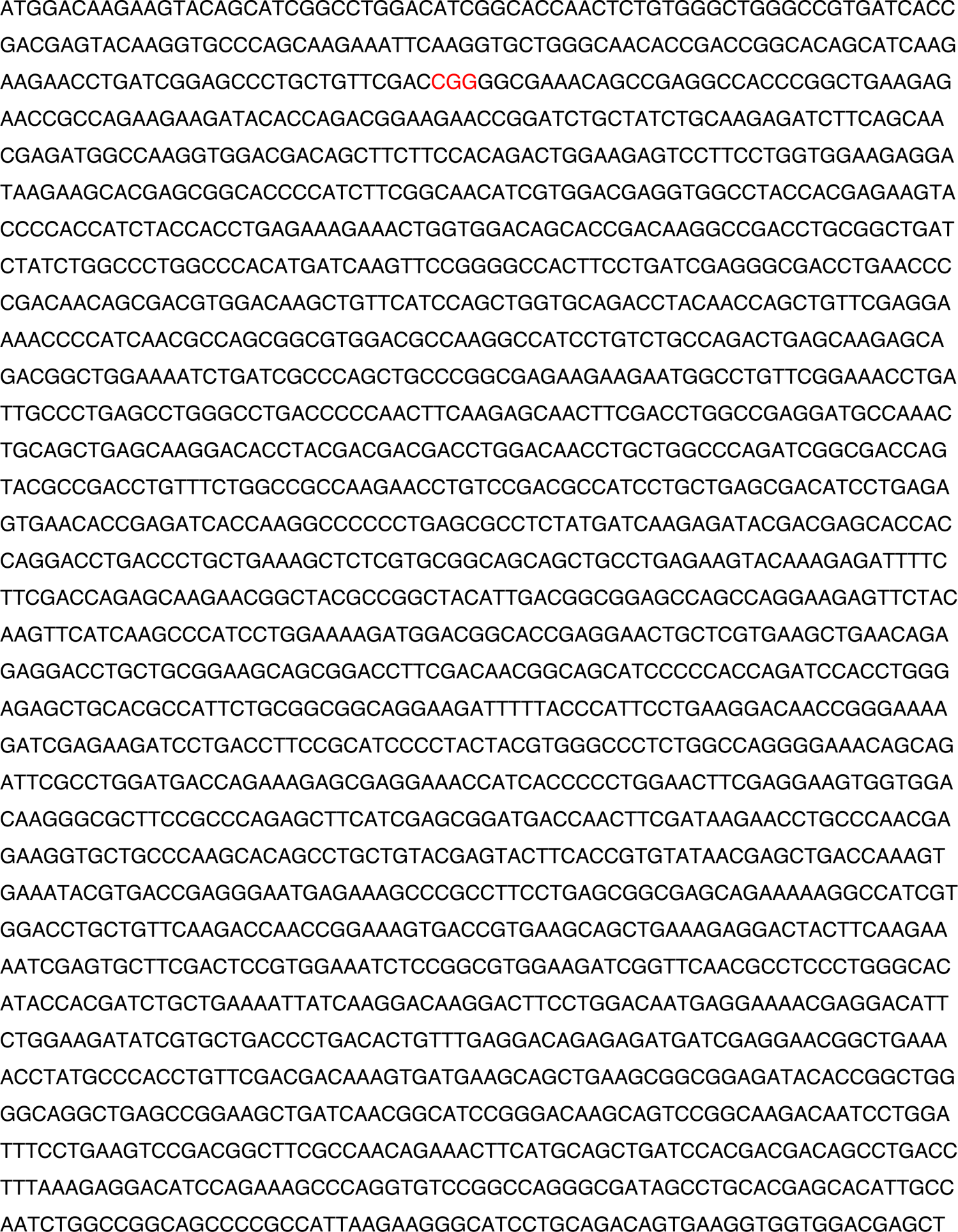
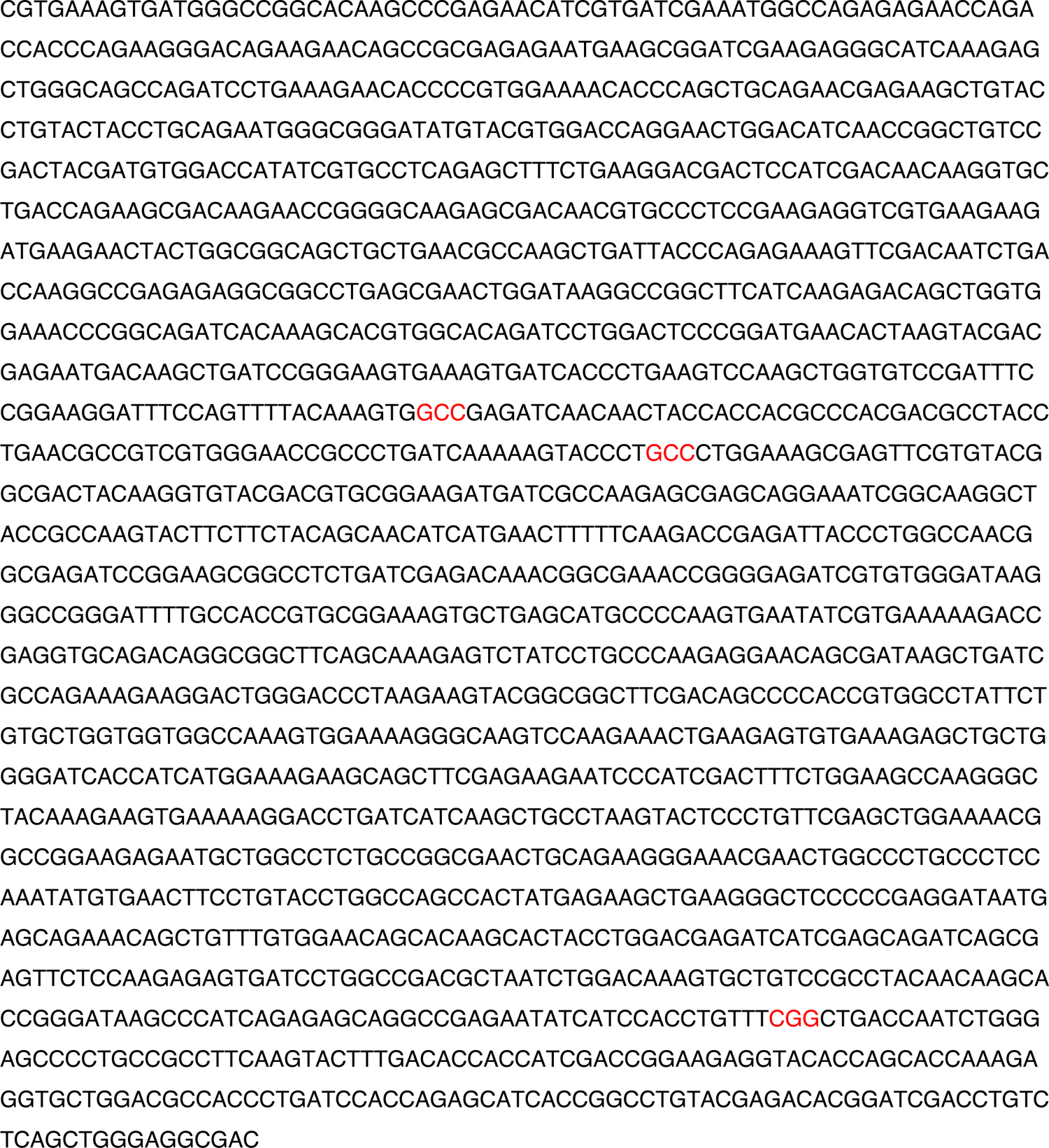

#### gRNA cloning backbone, with U6 promoter in green and scaffold in blue

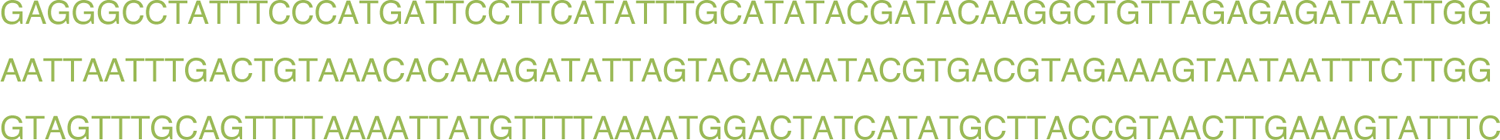
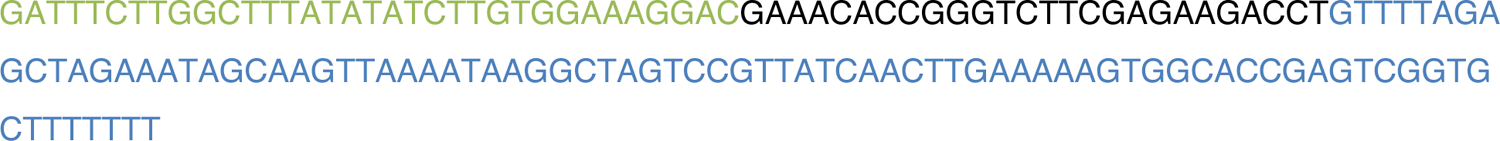

